# pH-dependent allosteric remodeling of a bacterial riboswitch couples alkaline activation to metal sensing

**DOI:** 10.64898/2026.02.18.706685

**Authors:** Danea Palmer, Adrien Chauvier, Tomás F. D. Silva, Avery Ontiveros, Giovanni Bussi, Nils G. Walter, Tatiana V. Mishanina

## Abstract

The widespread *yybP-ykoY* riboswitches control bacterial manganese (Mn) homeostasis by activating exporter expression in response to intracellular Mn^2+^ levels. The *E. coli alx* riboswitch distinctively couples Mn^2+^ sensing to cytoplasmic alkalinity, but the mechanism is unknown. We show that pH tunes the *alx* aptamer’s conformational sampling to modulate Mn^2+^ sensitivity. Single-molecule FRET reveals that Mn^2+^ stabilizes a docked three-way-junction conformation, and alkaline pH shifts this equilibrium to sensitize metal-dependent folding. Molecular dynamics simulations identify a loop whose low-pH-induced base pairing perturbs the adjacent helix, predicted to allosterically disrupt the Mn^2+^-binding state. *In vivo* reporters indicate that both this loop and the Mn^2+^-binding core are required for optimal pH-dependent translational activation: replacing the core with the non-pH-responsive *mntP* sequence abolishes activation. These results define how RNA allosterically integrates orthogonal metal and proton cues to enable combinatorial environmental sensing during alkaline stress.

## INTRODUCTION

Bacteria inhabit diverse, fluctuating environments and have evolved rapid responses to numerous, often overlapping stresses, such as alkaline pH and changes in micronutrient metal availability^1,2^. Transition metals are essential cofactors for nearly half of all cellular enzymes, but an excess of these metal ions can compromise cell survival^3,4^. Riboswitches are one gene regulatory mechanism by which bacteria achieve metal ion homeostasis^5,6^. These structured non-coding RNA elements are embedded within the 5ʹ untranslated regions (5ʹ-UTR) of messenger RNAs (mRNAs). Upon binding their cognate ligands with exquisite selectivity, riboswitches undergo conformational changes that promote or inhibit expression of a downstream gene^7^, thereby acting as allosteric biomolecular sensors of the cellular state. Each riboswitch comprises a ligand-binding aptamer domain and an expression platform that either controls transcription termination or affects translation initiation by dictating mRNA ribosome binding site (RBS) accessibility^7^. Metal ion-binding riboswitches typically modulate the transcription or translation of genes encoding metal ion transport proteins^6^. A variety of metal ions serve as ligands, including magnesium (Mg)^8,9^, manganese (Mn)^10,11^, cobalt (Co)^12^, nickel (Ni)^12^, lithium (Li)^13^, and sodium (Na)^14^. Among these, Mn^2+^-sensing motifs constitute the largest class of metal-ion sensing riboswitches and belong to the *yybP-ykoY* family that is broadly distributed among bacteria^5,15^.

One unique member of the *yybP-ykoY* family found in *Escherichia coli* (*E. coli*), *alx,* was originally identified as a “pH-responsive element” that activated translation of the downstream gene at alkaline pH^16^. Later, *alx* was shown to be a Mn^2+^-sensing riboswitch^10^, and the protein product of the downstream gene was recently characterized as an Mn^2+^ exporter^17^. *In vivo* gene reporters showed that, while Mn^2+^ supplementation alone boosts *alx* translation ∼20-fold, translation increases ∼70-fold when *E. coli* are cultured at alkaline pH^17^, demonstrating the remarkable ability of the *alx* riboswitch to incorporate two distinct inputs (metal ion and proton concentration) into a single gene expression output. In contrast, a second member of the *yybP-ykoY* family also found in *E. coli* (*mntP*) likewise upregulates translation of an Mn^2+^ exporter as a function of Mn^2+^ concentration, but shows no translational response to alkaline pH^17^.

The structures of several representative *yybP-ykoY* aptamers have been characterized, with riboswitches from *L. lactis, X. oryzae,* and *E. coli* (*mntP*) assuming a four-way junction (4WJ) architecture^11,18^. The 4WJ is comprised of four helices (P1-P4), which organize into two helical legs that dock to stabilize the Mn^2+^-binding site^11,18^. In contrast, the *alx* aptamer assumes a three-way junction (3WJ) architecture with a “capping loop”, L2, replacing the 4WJ P2 helix^19^ (Fig. 1a). Despite this difference, both the 3WJ and 4WJ aptamers share a structurally conserved metal-ion binding core. This core is composed of a conserved loop L1 in the “right leg” of the aptamer and a variable loop L3 in the “left leg” that juxtapose to coordinate two metal ions (Fig. 1a)^11^. One metal-ion binding site is occupied by a Mg^2+^ ion (M_A_) – the dominant, millimolar cellular divalent metal ion – whereas the other has micromolar affinity for Mn^2+^ (M_B_), specified by a direct contact with N7 of an absolutely conserved adenosine in L3 (A37 in *alx*; Fig. 1a)^11,19,20^.

**Fig. 1:**
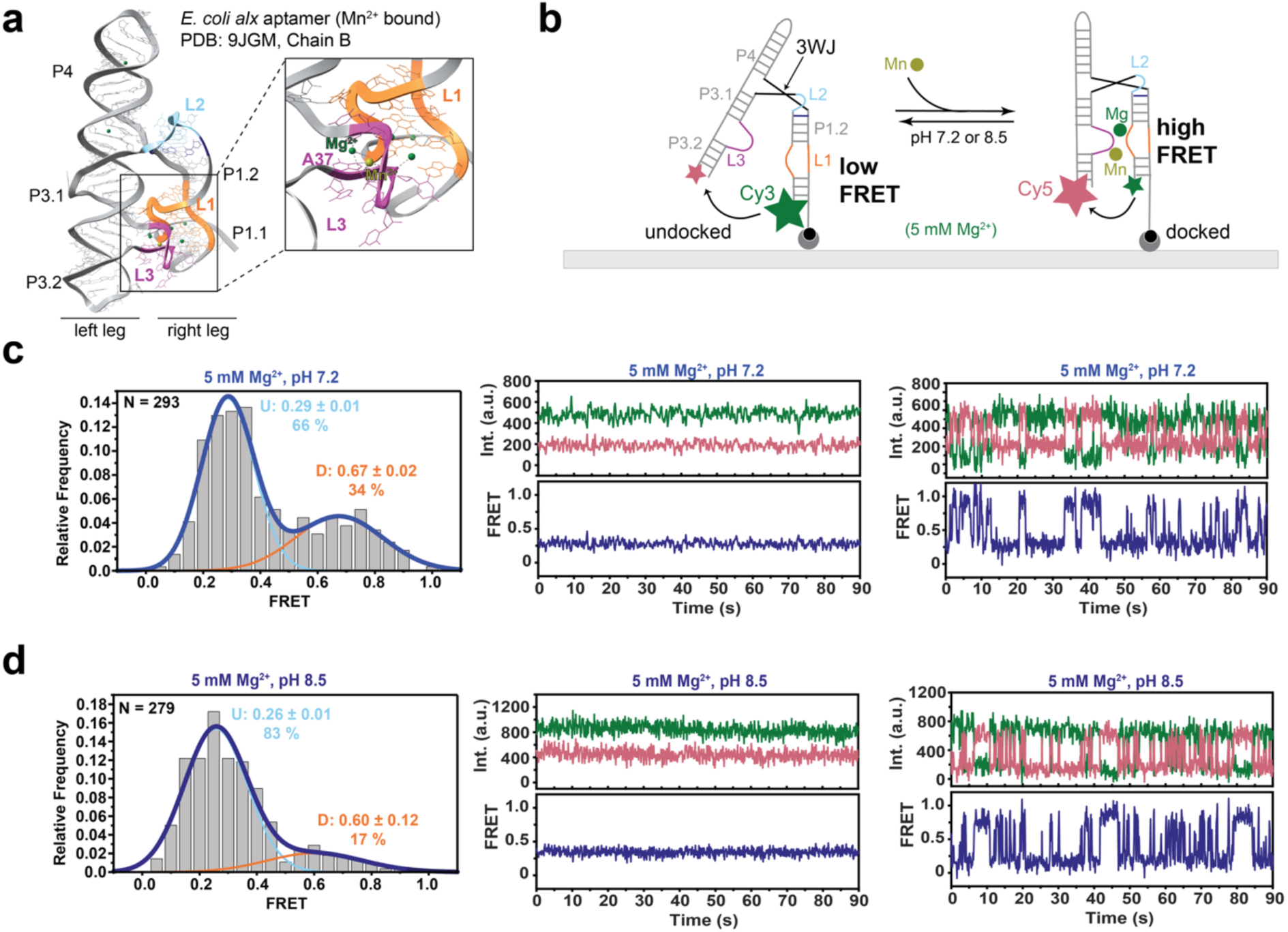
*alx* aptamer architecture and dynamics captured by smFRET at neutral and alkaline pH. **a,** *alx* aptamer crystal structure with a zoomed-in panel for the Mn^2+^ binding site (PDB: 9JGM, Chain D). **b,** Experimental setup for smFRET experiments. **c,d,** Histograms and representative traces showing the fraction of low and high FRET molecules at **(c)** pH 7.2 and **(d)** pH 8.5.

Previous <u>s</u>ingle-<u>m</u>olecule <u>F</u>örster <u>R</u>esonance <u>E</u>nergy <u>T</u>ransfer (smFRET) studies of 4WJ *yybP-ykoY* aptamers from *X. oryzae* and *L. lactis* demonstrated that, at physiological Mg^2+^ concentration, the RNA transiently samples the docked state, but becomes stably docked only in the presence of Mn^2+^^18,21,22^. Consistent with this behavior, molecular dynamics (MD) simulations indicated that Mn^2+^ binding is required to stabilize base stacking in the L3 loop^18,23^. A recent co-transcriptional *in vitro* folding analysis of the *E. coli mntP* and *alx* riboswitches found that alkaline pH alone does not appreciably alter overall folding of the *alx* riboswitch aptamer or its expression platform, but instead appears to modulate Mn^2+^ binding to the 3WJ^24^. *In vivo* translation reporter assays further showed that an A37U mutation of the Mn^2+^-coordinating adenosine – a mutation that abolishes Mn^2+^-dependent docking in the *X. oryzae* 4WJ aptamer^18^ – abrogates pH-dependent translation activation of the 3WJ *alx* aptamer^24^, suggesting that coupled pH and Mn^2+^ sensing may trigger exporter gene expression before Mn^2+^ toxicity rises under alkaline stress^24^. Notably, both *alx* and *mntP* aptamers dynamically sample ambient Mn^2+^ during co-transcriptional folding^24^. However, the molecular mechanism underlying pH-adaptive tuning of ligand binding remains unresolved; defining it should reveal principles by which RNAs can reweight their folding ensembles to allosterically integrate orthogonal chemical inputs, with implications for natural regulatory logic and the engineering of multi-input RNA sensors.

To address this gap, we used smFRET to resolve the real-time tertiary structure dynamics of the *alx* 3WJ riboswitch aptamer and found that pH modulates both aptamer docking and Mn^2+^ binding. In the absence of Mn^2+^, the *alx* aptamer exhibits low occupancy of a stably docked conformation at neutral pH, which is lost under alkaline conditions as the RNA remains predominantly undocked. By contrast, at sub-saturating Mn^2+^, alkaline pH more effectively promotes the stably docked state than neutral pH, thereby priming the aptamer for increased Mn^2+^ sensitivity in alkaline environments. MD simulations identify a pH-dependent nucleobase protonation event in the flexible L2 capping loop – unique to the 3WJ aptamer – that reorients the loop and provides a structural basis for pH-dependent docking and Mn^2+^ sensitization. Consistent with this model, translation reporter assays show that the resulting alternative base pairing in L2 is required for alkaline activation *in vivo*. Moreover, *in vivo* reporter analyses establish that the native *alx* L3 docking loop within the Mn^2+^-binding core is also required for pH-dependent Mn^2+^ responsiveness, supporting long-range allosteric coupling. Together, these results define a multipartite mechanism by which pH tunes riboswitch conformational dynamics to integrate metal and proton cues within a single aptamer and mitigate cellular Mn^2+^ toxicity under alkaline stress.

## RESULTS

### In the absence of Mn^2+^, alkaline pH reduces aptamer docking

To assess the 3WJ *alx* aptamer global dynamics, we used smFRET to monitor aptamer docking in real time under alkaline pH and elevated Mn^2+^ conditions. A FRET fluorophore pair (Cy3 and Cy5) was positioned on each leg of the riboswitch aptamer, and the relative fluorophore intensities were used to identify the low-FRET (undocked) and high-FRET (docked) states (Fig. 1b and Extended Data Fig. 1). smFRET traces at near-physiological Mg^2+^ concentration (5 mM) in the absence of Mn^2+^ revealed a highly dynamic ensemble of RNA molecules at both pH 7.2 (neutral) and pH 8.5 (alkaline). At pH 7.2, 66% of imaged aptamer molecules assumed the undocked conformation (Fig. 1c) compared to 83% at pH 8.5 (Fig. 1d). The mean low-FRET values are similar at both pHs (0.29 at pH 7.2 versus 0.26 at pH 8.5), indicating that the RNA samples very similar low-FRET states regardless of pH (Fig. 1c,d). Under all conditions, many traces captured transitions between low and high FRET states (Fig. 1c,d), consistent with smFRET data for the 4WJ Mn^2+^-sensing aptamers at physiological Mg^2+^ conditions^18,21,22^.

To further evaluate the docking dynamics, we employed transition occupancy density plots (TODP)^25^, ensemble heatmaps representing each molecule’s behavior: molecules undergoing at least one transition between FRET states appear off-diagonal, whereas molecules that never transition (i.e., stably remain in one state) are found on the diagonal (see Methods). At neutral pH (pH 7.2) in the absence of Mn^2+^ (with 5 mM Mg^2+^), the TODP shows that ∼17% of aptamer molecules are stably docked (SD), without noticeable undocking (Fig. 2a and Supplementary Table 5). In contrast, the 4WJ aptamers from *L. lactis* and *X. oryzae* show no significant SD complex unless Mn^2+^ is present^18,22^. In addition, a significant fraction of the *alx* traces is dynamic and exhibits double-exponential FRET state interconversion kinetics with k_dock,fast_ ∼2.0 s^-1^ (73%) and k_dock,slow_ ∼0.26 s^-1^ (27%), as well as k_undock,fast_ ∼3.89 s^-1^ (63%) and k_undock,slow_ ∼0.71 s^-1^ (37%; Fig. 2e). Accordingly, ∼30% of aptamer molecules populate a dynamically undocked (DU) state, whereas ∼11% occupy a dynamically docked (DD) state. The remaining 42% are stably undocked (SU) (Fig. 2a and Supplementary Table 5). Overall, the weighted-mean docking rate constant (k_dock,overall_ ∼1.53 s^-1^) is slower than the corresponding undocking rate constant (k_undock,overall_ ∼2.71 s^-1^; Supplementary Table 6), favoring the undocked state.

**Fig. 2:**
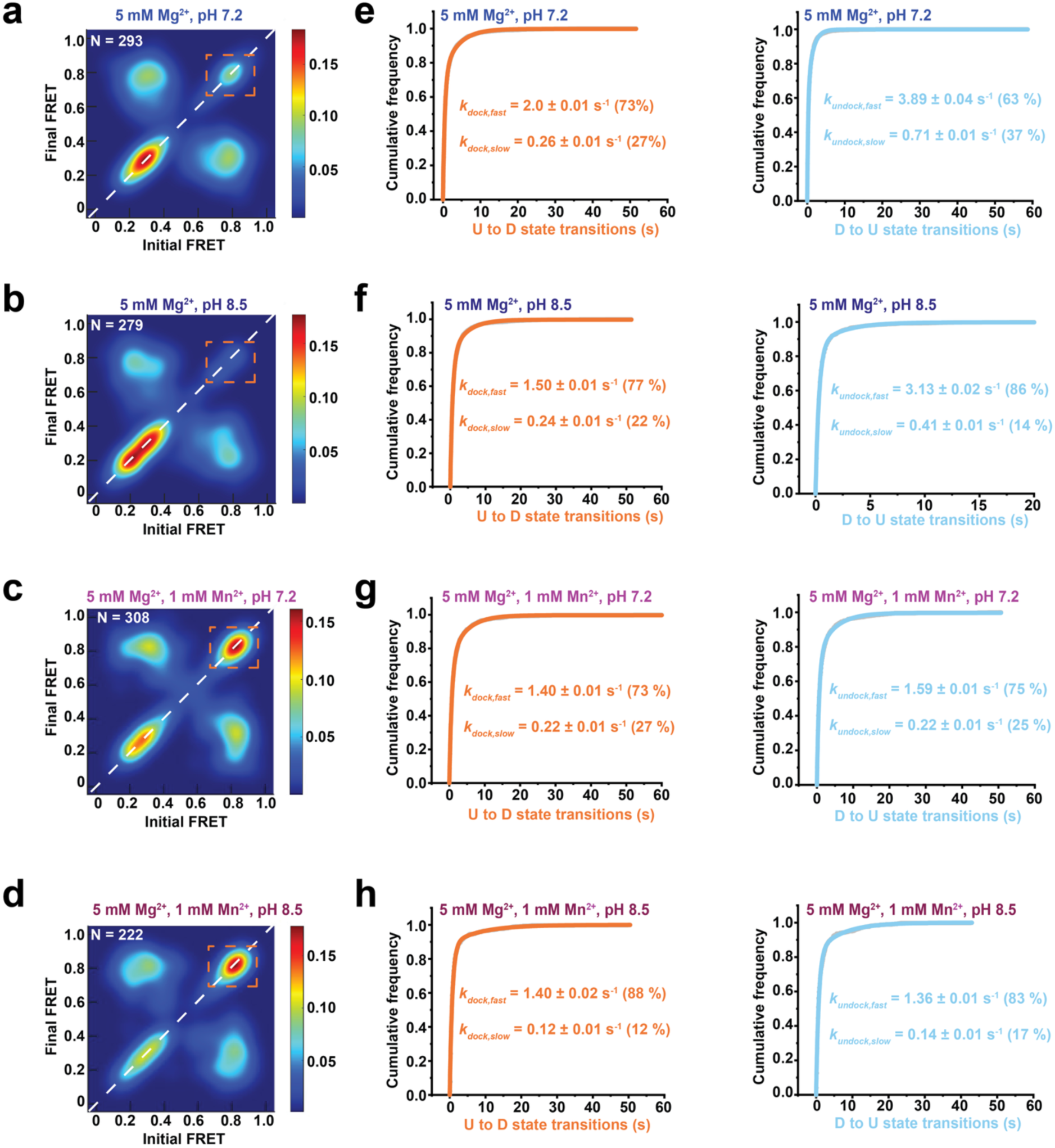
Saturating Mn^2+^ ligand stabilizes docked *alx* aptamer and impacts docking kinetics. **a-d,** Transition occupancy density plot (TODP) showing initial FRET vs final FRET. Traces on the diagonal represent static molecules and traces off the diagonal represent dynamic molecules, with the stable docked (SD) conformation highlighted with the dashed box. TODPs for **(a)** pH 7.2 with 5 mM Mg^2+^, **(b)** pH 8.5 with 5 mM Mg^2+^, **(c)** pH 7.2 with 5 mM Mg^2+^ and 1 mM Mn^2+^, **(d)** pH 8.5 with 5 mM Mg^2+^ and 1 mM Mn^2+^. **e-h,** Kinetics for docking (U to D state transitions) and undocking (D to U state transitions) based on the cumulative dwell times in each state fit with double-exponential functions for **(e)** pH 7.2 with 5 mM Mg^2+^, **(f)** pH 8.5 with 5 mM Mg^2+^, **(g)** pH 7.2 with 5 mM Mg^2+^ and 1 mM Mn^2+^, **(h)** pH 8.5 with 5 mM Mg^2+^ and 1 mM Mn^2+^.

Under alkaline conditions (pH 8.5) in the absence of Mn^2+^, only ∼9% and ∼6% of traces reside in the SD and DD conformations, respectively (Fig. 2b and Supplementary Table 5). This pronounced loss of docking ability is consistent with prior smFRET analyses of the *L. lactis* and *X. oryzae* 4WJ Mn^2+^-sensing aptamers^18,22^. The docking kinetics of dynamic traces is again well described by a double-exponential model, with k_dock,fast_ ∼1.5 s^-1^ (77%) and k_dock,slow_ ∼0.24 s^-1^ (22%; Fig. 2f), whereas undocking occurs with k_undock,fast_ ∼3.13 s^-1^ (86%) and k_undock,slow_ ∼0.41 s^-1^ (14%; Fig. 2f). As at neutral pH, the weighted-mean docking rate constant k_dock,overall_ ∼1.21 s^-1^ is slower than undocking at k_undock,overall_ ∼2.75 s^-1^ (Supplementary Table 6). Relative to docking at neutral pH, k_dock,overall_ decreases, whereas k_undock,overall_ remains largely unchanged. Consistent with this shift, a slightly higher ∼35% of traces populate the DU state, and only ∼6% remain in the DD state―nearly a two-fold reduction compared to pH 7.2 (Fig. 2b and Supplementary Table 5). Notably, 50% of *alx* aptamer molecules reside in the SU state at alkaline pH, exceeding both the neutral pH fraction (42%) and the SU populations previously reported for the 4WJ *L. lactis* and *X. oryzae* aptamers^18,22^. Together, these data indicate that, in the absence of Mn^2+^, alkaline pH destabilizes docked conformations and slows docking transitions, biasing the 3WJ aptamer toward a more undocked ensemble that may prime it for increased Mn^2+^ sensitivity upon ligand availability.

### Saturating Mn^2+^ promotes docking independent of pH by slowing undocking

Despite the pH-dependent differences in the docking equilibrium of the apo *alx* aptamer, addition of 1 mM Mn^2+^ (orders of magnitude higher than physiological ∼20 µM^17^) to the aptamer produced remarkably similar smFRET traces at both neutral and alkaline pH, but consistently favored docking. In accordance with 4WJ Mn^2+^-sensing aptamers, Mn^2+^ binding to the 3WJ *alx* aptamer stabilizes the SD conformation, with ∼29% and ∼31% SD molecules at pH 7.2 and pH 8.5, respectively (Fig. 2c,d and Supplementary Table 5). Additionally, the mean high FRET value upon Mn^2+^ binding at both pHs is 0.76, indicating that the RNA is sampling the same docked conformation (Extended Data Fig. 2a,b). Despite the presence of saturating Mn^2+^, ∼30% of molecules remain SU at both pHs. This is drastically different from the 4WJ *L. lactis* and *X. oryzae* aptamers, which dock completely with saturating Mn^2+18,22^. At both pHs, around 40% of *alx* traces are dynamic in the absence and presence of Mn^2+^, but significantly more of them are DD in the presence of 1 mM Mn^2+^ (∼20% versus ∼6% in the absence of ligand). The increase in DD traces upon Mn^2+^ binding correlates with a decrease in DU traces (Supplementary Table 5), serving as additional evidence that Mn^2+^ promotes the docked conformation. These dynamic molecules again exhibit double-exponential kinetics, with k_dock,overall_ ∼1.08 s^-1^ and k_undock,overall_ ∼1.25 s^-1^ at pH 7.2 and k_dock,overall_ ∼1.25 s^-1^ and k_undock,overall_ ∼1.15 s^-1^ at pH 8.5 (Fig. 2g,h and Supplementary Table 6). Rather than speeding *alx* aptamer docking, ligand binding slows undocking by >2-fold regardless of pH. Similarly, Mn^2+^ binding slows aptamer undocking for the 4WJ *L. lactis* and *X. oryzae* aptamers, indicative of a conserved mechanism for the effect of Mn^2+^ binding in the *yybP-ykoY* riboswitch family^18,21,22^. Overall, regardless of pH, ligand binding shifts the *alx* aptamer conformational equilibrium toward the docked state, primarily by slowing undocking.

### Alkaline pH sensitizes the 3WJ *alx* aptamer to Mn^2+^ *in vitro*

Because pH changes did not alter *alx* aptamer dynamics at saturating ligand, we hypothesized that alkaline pH instead biases the *alx* aptamer toward an open conformation that increases sensitivity to Mn^2+^. To test this idea, we recorded smFRET traces at pH 7.2 and pH 8.5 in the presence of a low Mn^2+^ concentration (1 µM; Fig. 3a,b). At pH 7.2, a 72% population resides in the undocked state with a low FRET value of 0.28 (Fig. 3a), comparable with the no-Mn^2+^ condition. Accordingly, the TODPs showed largely unchanged dynamics upon addition of 1 µM Mn^2+^ at pH 7.2 (Fig. 2a and Fig. 3c), and undocking kinetics were essentially identical to no Mn^2+^ (Extended Data Fig. 3a), indicating that 1 µM Mn^2+^ is insufficient to perturb *alx* aptamer dynamics at neutral pH. In contrast, at pH 8.5, 1 µM Mn^2+^ significantly shifted the population toward the docked conformation (37% docked; Fig. 3b) compared to 17% in the absence of Mn^2+^ (Fig. 1d). Moreover, the conformational dynamics at pH 8.5 with 1 µM Mn^2+^ closely resembled those at saturating 1 mM Mn^2+^ (Fig. 2b and Fig. 3d), with ∼26% SD and ∼18% DD traces (Fig. 3d and Supplementary Table 5). Fits yielded k_dock,overall_ ∼1.20 s^-1^ and k_undock,overall_ ∼1.89 s^-1^ at pH 8.5 with 1 µM Mn^2+^ (Extended Data Fig. 3b and Supplementary Table 6). Although 1 µM Mn^2+^ slows undocking less than 1 mM Mn^2+^, the reduction remains substantial. Together, these data indicate that alkaline pH uniquely enables the *alx* aptamer to bind and respond to low ligand concentrations to trigger an effective gene regulatory response.

**Fig. 3:**
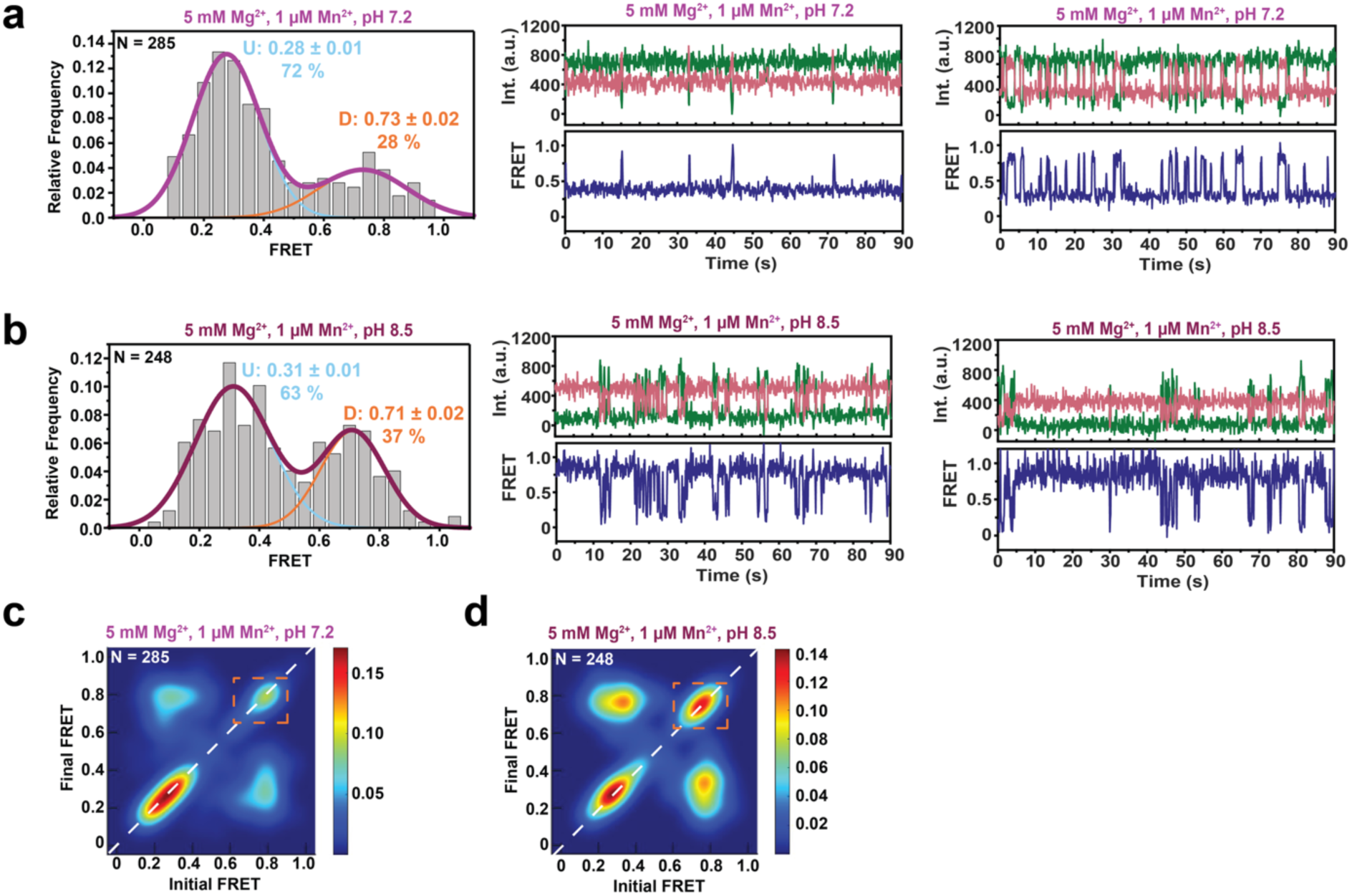
Low Mn^2+^ concentration uniquely promotes docked *alx* aptamer at alkaline pH. **a,b,** Histograms and representative traces showing the fraction of low and high FRET molecules with 1 µM Mn^2+^ at **(a)** pH 7.2 and **(b)** pH 8.5. Histograms display the Gaussian peaks for low FRET, high FRET, and cumulative fit with the percentage of each FRET state and the number of molecules (N) analyzed. **c,d,** TODPs showing the initial FRET vs final FRET with 1 µM Mn^2+^ at **(c)** pH 7.2 and **(d)** pH 8.5. Traces on the diagonal represent static molecules and traces off the diagonal represent dynamic molecules, with the stable docked (SD) conformation emphasized in the dashed box.

### The L2 capping loop couples pH sensing to Mn^2+^-dependent gene activation

In contrast to the canonical 4WJ *yybP-ykoY* aptamer architecture, *alx* adopts a 3WJ fold in which the P2 helix is replaced by a single-stranded loop (Fig. 4a)^19^. In the *alx* aptamer crystal structure, L2 is stabilized by two P1 base pairs, U18·A114 and C19·G23, forming a 3-nucleotide (nt) “capping loop”^19^. Because L2 is unique to *alx* and located at the junction between the two docking legs, we hypothesized that it contributes to pH-dependent Mn^2+^ sensing.

**Fig. 4:**
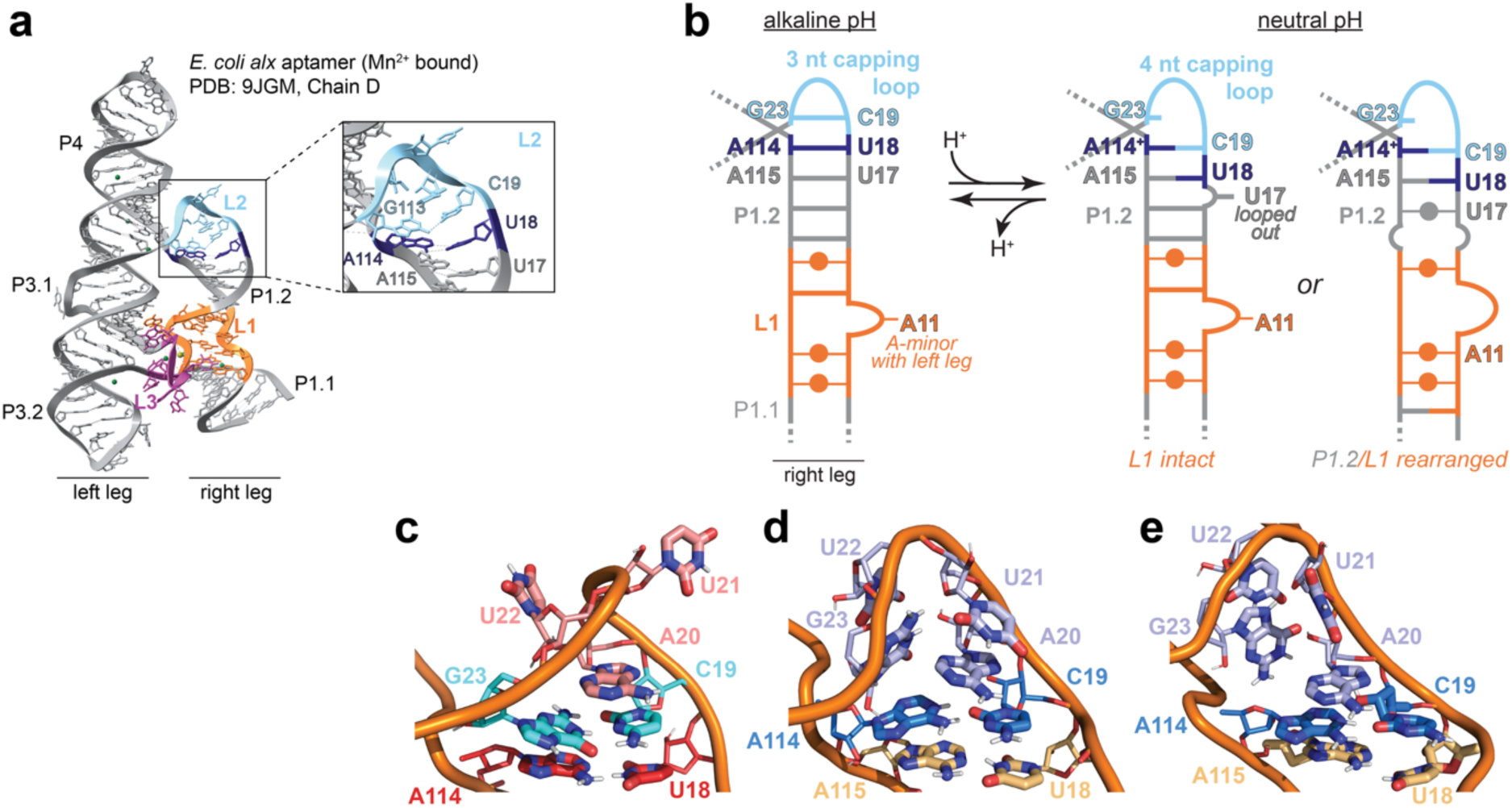
L2 capping loop length depends on A114 protonation and stability of key base pairs. **a,** *alx* aptamer crystal structure with a zoomed-in panel for L2, with key nucleotides labeled (PDB: 9JGM, Chain D). **b,** Cartoon representation of proposed alternative base pairing at alkaline and neutral pH, with key nucleotides labeled. **c-e,** Cartoon representations of L2 capping loop with the **(c)** native, **(d)** wobble, and **(e)** an alternative meta-stable state, generated with PyMOL^98^. In panel (c), the 3-nt L2 loop formed by U22, U21, and A20 (pink) requires stable U18·A114 (red) and C19·G23 (cyan) base pairs. In panels (d) and (e), the 4-nt L2 loop (light purple) requires the U18·A115 (beige) base pair and, at least, the O2·N1H^+^ hydrogen bond formed within the C19·A114^+^ (blue) base pair.

To evaluate this idea, we reanalyzed previously reported co-transcriptional RNA structure probing data for the *alx* riboswitch collected at pH 7.2 or pH 8.5 with or without 1 mM Mn^2+24^, matching the conditions used here. Probing was performed with DMS, which under these conditions primarily reports on A(N1) and C(N3) accessibility^26^; thus our analysis focuses on A and C nucleobases. A114 and A115 shows reduced DMS reactivity at pH 8.5 relative to pH 7.2, independent of Mn^2+^ (Extended Data Fig. 4a,b). C19 also exhibits a marked decrease in reactivity at pH 8.5, but only beginning at transcript length of 127 nt, after the transcription of the A114 and A115 base-pairing partners (Extended Data Fig. 4c). These findings indicate that the structure of L2 is more stabilized at alkaline pH, whereas its higher DMS reactivity at pH 7.2 is consistent with greater flexibility. As expected for a loop nucleotide, A20—predicted to be highly flexible— remains highly reactive at both pH values throughout aptamer synthesis (Extended Data Fig. 4d).

These pH-dependent changes led us to hypothesize that pH may be altering the length of the L2 capping loop to modulate docking in the 3WJ *alx* aptamer: protonation of A114 could enable a 4-nt RNA via formation of a C19·A114^+^ wobble pair, whereas A114 deprotonation at alkaline pH would favor a strained 3-nt capping loop stabilized by canonical base pairing (Fig. 4b). Formation of the non-canonical C19·A114^+^ interaction, together with the disruption of the adjacent U17·A115 base pair in the adjacent helix P1.2, could account for the higher DMS reactivity of these nucleotides at pH 7.2 (Fig. 4b and Extended Data Fig. 4a-c). Conversely, strain associated with a 3-nt loop at alkaline pH may contribute to the static undocked conformation observed by smFRET at pH 8.5 (Fig. 2b). Although adenosines have a solution pK_a_ of ∼3.5, local RNA environments can substantially shift pK_a_ values^27^, and C·A^+^ wobble pairs can be functionally essential in cells^28,29^.

To test whether C19·A114^+^ pairing is possible under our experimental conditions, we performed molecular dynamics (MD) simulations based on the published *alx* aptamer crystal structure (PDB ID: 9JGM)^19^, with either neutral or protonated A114. The neutral A114 maintained stable base pairing over the 200 ns (Fig. 4c). Constant-pH MD (CpHMD) simulations of the native structure yielded a neutral A114 across the tested pH range (3.0-7.0), consistent with stabilization of the U18·A114 Watson-Crick base pair^30,31^. In contrast, during simulations initiated with a protonated adenosine, the U18·A114 Watson-Crick pair ruptured; however, A114^+^ and C19 remained separated (>3 Å), preventing spontaneous formation of an C19·A114^+^ wobble (Extended Data Fig. 5a,b). Forcing an C19·A114^+^ wobble enabled hydrogen bonding, but the interaction was unstable (Extended Data Fig. 5b).

Because flanking base pairs can stabilize wobble interactions^27^, we next considered formation of an additional Watson-Crick pair (U18·A115) involving A115 from the adjacent helix P1.2. Enforcing simultaneous C19·A114^+^ and U18·A115 pairing, guided by a model duplex containing a wobble base pair flanked by a A·U pair^32^, produced a metastable structure (Fig. 4d and Extended Data Fig. 5b and 6b), supporting the potential of C19·A114^+^ formation at neutral pH with an accompanying rearrangement of helix P1.2 in the aptamer’s “right” leg. Notably, the titrating proton (N1H of A114) could also be stabilized by acceptors beyond the N3 of C19 (Fig. 4e and Extended Data Fig. 7a). Across simulations, structural analysis of A114 protonation supports a 4-nt L2 capping loop, with C19·A114^+^ stabilized by at least one hydrogen bond involving the titrating proton (Fig. 5a and Extended Data Fig. 7b), whereas the absence of A114 protonation at alkaline pH favors the formation of a 3-nt L2 loop (Extended Data Fig. 8). To estimate the pK_a_ for the A114 N1H, we restrained the fully formed C19·A114^+^ wobble pair and performed constant pH simulations^33^, yielding a microscopic pK_a_ of ∼12 for this conformation (Fig. 5b). Because the macroscopic pK_a_ reflects a weighted average across conformations, and because the C19·A114^+^ wobble was only partially stable in our simulations, the effective pK_a_ could be closer to 8.5, consistent with the strong *alx* induction *in vivo*^17^ and pronounced smFRET changes *in vitro* (Fig. 3d).

**Fig. 5:**
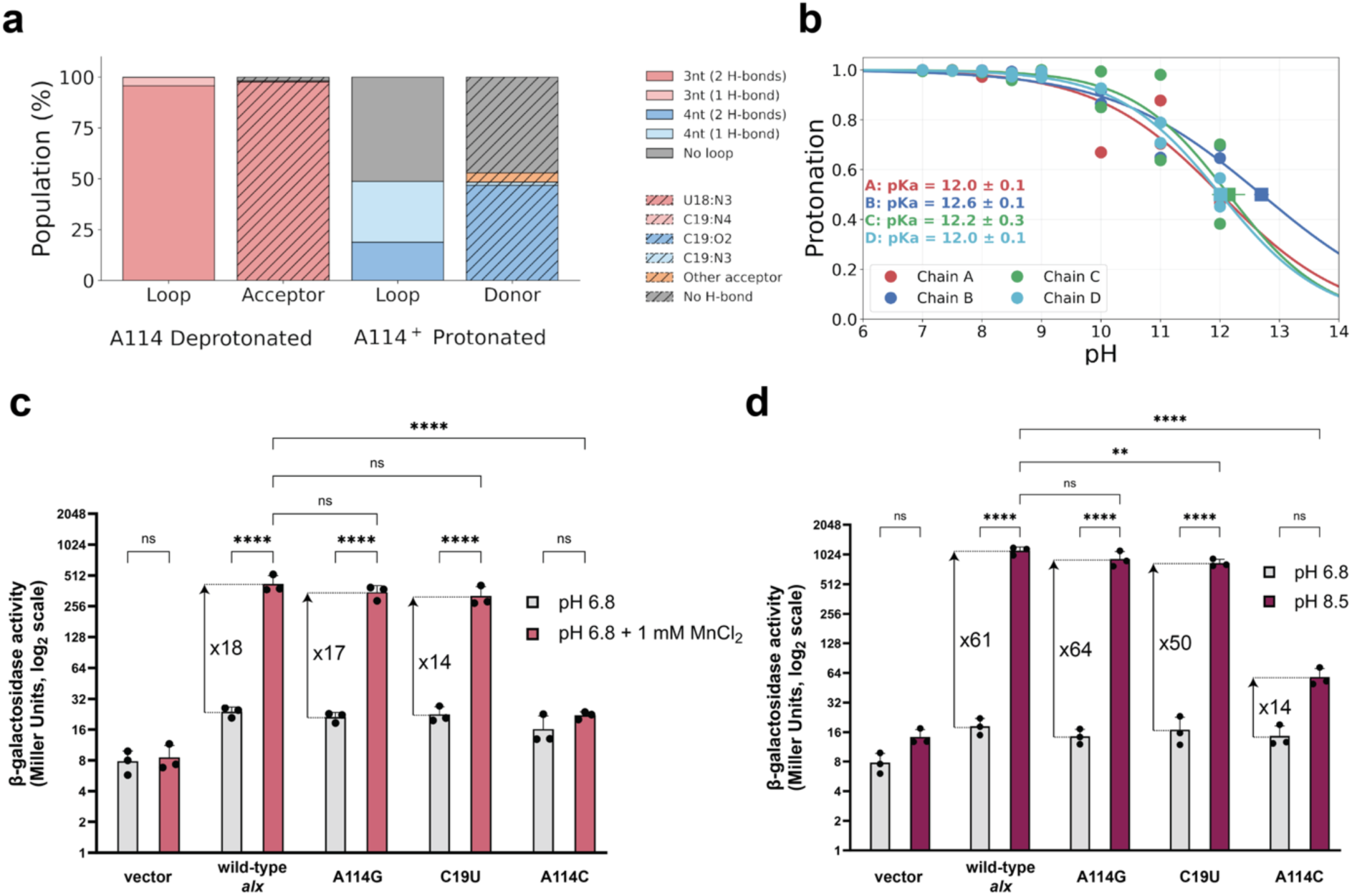
L2 capping loop mutations reduce pH-dependent *alx* translation activation. **a,** Predicted relative population (%) of loop-length states and specific hydrogen-bond contacts shown by the hydrogen-bonding role (acceptor or donor) of A114 dependent on the protonation state (deprotonated/protonated) and grouped by the number of hydrogen-bonds. **b,** Titration curves and microscopic pKa values of A114 obtained from MD simulations of the wobble conformation state at constant pH, for each crystallographic chain (A-D). Mean protonation values from independent replicates were fit to the Henderson–Hasselbalch equation. Upshifted pKa values highlight the proton chemical stability even in highly basic environments, as a result of the specific hydrogen-bond network. **c,d,** Translational response for L2 mutants compared to wild-type *alx* for growth media **(c)** supplemented with Mn^2+^ and **(d)** buffered at alkaline pH. Statistical analysis done using a two-way ANOVA with Tukey post-hoc test for multiple comparisons to assess significance (^ns^ P>0.05, * P≤0.05, ** P≤0.01, *** P≤0.001, **** P≤0.0001).

**Fig. 6:**
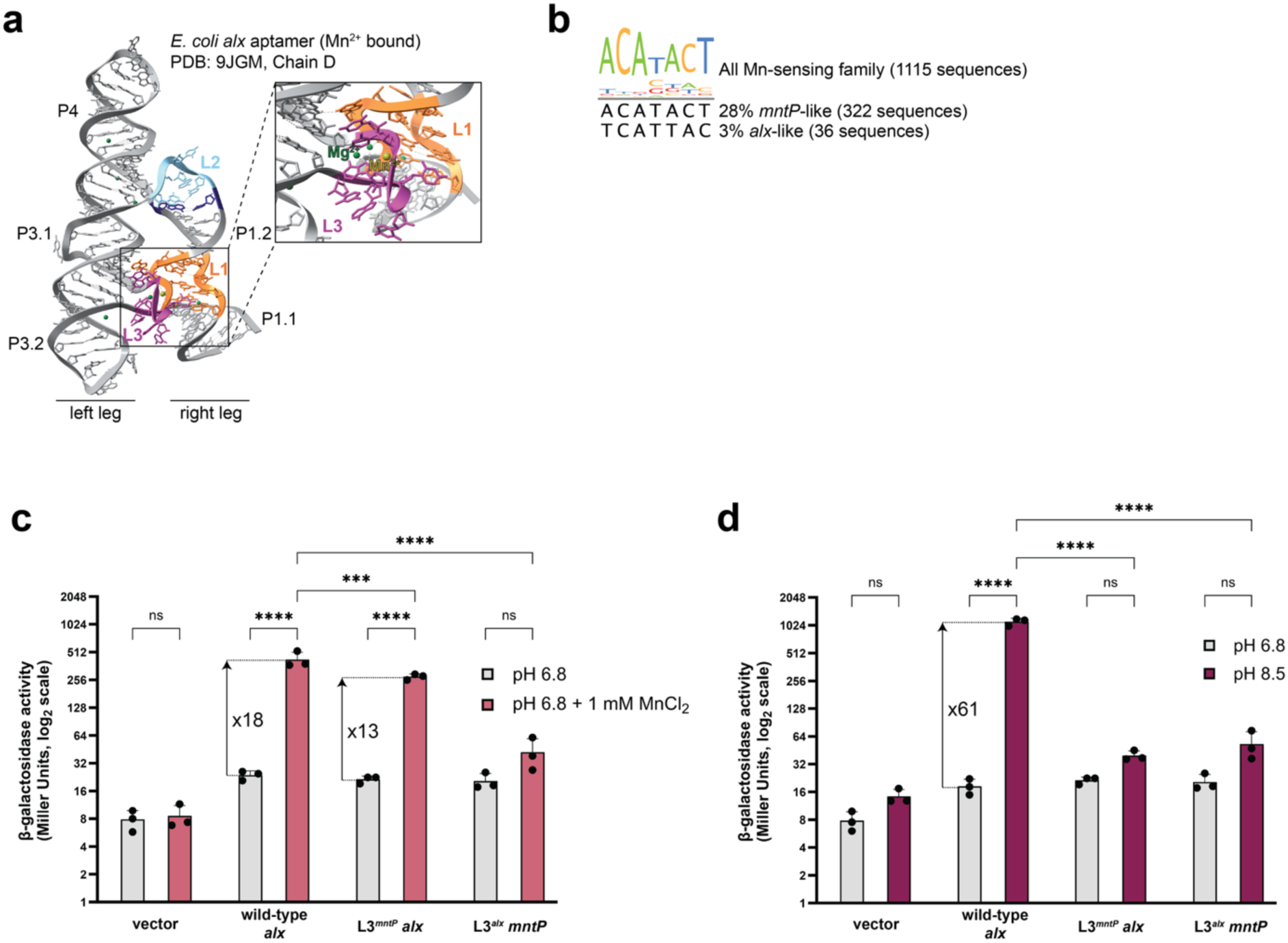
Native *alx* L3 loop sequence is essential for pH-dependent translation activation. **a,** *alx* aptamer crystal structure with a zoomed-in panel for Mn^2+^ binding site, with key loops L1 and L3 labeled (PDB: 9JGM, Chain D). **b,** L3 consensus sequence from *yybP-ykoY* family members in Rfam database, with the percent of *mntP*-like and *alx*-like sequences displayed below the consensus sequence. **c,d,** Translational response for L3 loop swap (between *alx* and *mntP*) mutants compared to wild-type *alx* for growth media **(c)** supplemented with Mn^2+^ and **(d)** buffered at alkaline pH. Statistical analysis done using a two-way ANOVA with Tukey post-hoc test for multiple comparisons to assess significance (^ns^ P>0.05, * P≤0.05, ** P≤0.01, *** P≤0.001, **** P≤0.0001).

Guided by these simulations, we next tested the contribution of L2 to alkaline activation *in vivo*. We constructed *alx*-controlled translational *lacZ* gene reporters in single-copy plasmids bearing point mutations (A114G, C19U, and A114C) and quantified Mn^2+^- and pH-dependent expression using β-galactosidase assays, as previously described^17^. Two mutations, A114G and C19U, were designed to stabilize a 4-nt L2 capping loop, whereas A114C was expected to disrupt L2 base pairing. A114G and C19U did not significantly affect Mn^2+^ responsiveness compared to wild-type *alx* (Fig. 5c). At pH 8.5, A114G retained wild-type activation, while C19U reached 50-fold translation activation compared to 61-fold for wild-type *alx* (Fig. 5d). In contrast, A114C both abolished the Mn^2+^ response and strongly reduced pH-dependent translation activation (Fig. 5c,d). At a slightly less alkaline pH (pH 8.3), wild-type *alx* displayed 26-fold activation, while A114G and C19U were significantly reduced (12-fold and 9-fold, respectively; Extended Data Fig. 9), supporting an apparent pK_a_ near ∼8.5 for the pH-sensitive nucleoside(s) within the *alx* riboswitch. Together, our MD simulations and *in vivo* mutational analyses indicate that base pairing within the flexible L2 capping loop—likely involving pH-induced alternative pairing—is critical for pH- dependent translation activation, but becomes less relevant under increasingly alkaline conditions.

### The metal-binding L3 loop sequence is also required for the pH-dependent Mn^2+^ response

Because L2 capping loop mutations reduced—but did not abolish—pH-enhanced translation activation, we reasoned that additional riboswitch elements contribute to the full pH-dependent Mn^2+^ response of *alx*. Prior work showed that alkaline pH affects Mn^2+^ binding by *alx* and the pH-dependent gene expression response requires Mn^2+^ binding to the aptamer^24^. In *yybP-ykoY* riboswitches, the Mn^2+^-binding site forms between the conserved L1 loop and the variable L3 loop, which positions the conserved adenosine (A37 in *alx*) for direct coordination of Mn^2+^ (Fig. 6a)^11,20^. Across the family, L3 is ubiquitous: 28% of aptamers contain the *mntP* sequence (ACAUACU), whereas only 3% contain the *alx* sequence (UCAUUAC) (Fig. 6b); the remaining ∼70% differ from both by at least one nucleotide. Although the overall Mn^2+^-binding-site architecture appears similar across L3 variants (Extended Data Fig. 10a), we hypothesized that nucleotide identity within L3 modulates the stacking interactions that underlies pH-dependent Mn^2+^ binding. Specifically, we reasoned that after the riboswitch legs dock (tuned in part by the L2 capping loop), the L3 sequence further contributes to pH-dependent Mn^2+^ binding.

To test this notion, we swapped the L3 sequences between the pH-responsive *alx* aptamer and the non-pH responsive *mntP* aptamer^17^ in their respective translational reporter constructs and quantified output using *in vivo* β-galactosidase assays. The L3*^alx^ mntP* chimera completely lost the Mn^2+^ responsiveness and did not yield any gain in pH-dependent translation activation (Fig. 6c,d). Conversely, L3*^mntP^ alx* retained Mn^2+^ responsiveness, exhibiting a 13-fold induction at saturating Mn^2+^ compared with an 18-fold increase for wild-type *alx*, (Fig. 6c), but strikingly lost pH-dependent activation at pH 8.5 (Fig. 6d). These data demonstrate that the wild-type *alx* L3 sequence is essential for alkaline activation in cells.

To assess whether this alkaline stress response mechanism might be conserved across Mn^2+^-sensing riboswitches, we aligned all *yybP-ykoY* family members (1,115 sequences in Rfam)^34^, filtered for sequences containing the *alx* L3, and used RNAfold^35^ to identify aptamers predicted to adopt an *alx-*like 3WJ architecture, given the apparent importance of the L2 capping loop (Supplementary Table 7). Phylogenetic analysis showed that *yybP-ykoY* members with the *alx* L3 sequence are restricted to the *Bacillales* order, with 67% belonging in the *Bacillaceae* family (Extended Data Fig. 10b). Predicted *alx-*like 3WJ architectures are distributed more broadly across multiple families (Extended Data Fig. 10b). Notably, organisms containing both the *alx* L3 sequence and 3WJ architecture are found in soil or ocean environments and are capable of growth at alkaline pH^36–41^, with most organisms optimally growing at alkaline pH^36–40^. Together, our observations suggest that both the 3WJ global structure and the specific nucleotide context within the Mn^2+^-binding site are required for bacteria optimally adapted to alkaline stress^36–41^.

## DISCUSSION

Integrating smFRET, MD simulations, and *in vivo* gene reporters, we uncover how two distinct cellular cues—pH and Mn^2+^—are coupled to empower a rapid bacterial response to alkaline stress through riboswitch-mediated induction of a Mn^2+^ exporter. The 3WJ *alx* aptamer is strongly pH sensitive: alkaline pH favors an undocked (open) global architecture, whereas neutral pH supports more docked sampling, thereby priming the aptamer to specifically respond to low-µM Mn^2+^ under alkaline conditions. Our data further identify two key regions that together enable this dual-input behavior: the L2 capping loop at the junction between the docking legs and the L3 loop within the metal-binding core. In this way, *alx* provides an example of combinatorial gene regulation in bacteria in which riboswitch signal integration is key for cell survival in an alkaline habitat. Whereas eukaryotes often deploy large repertoires of transcription factors to integrate inputs^42^, bacteria frequently rely on such RNA-based integrative regulation to match metal-ion and metabolite availability to cellular demands^43,44^.

The pH-dependent control of Mn^2+^ export may be particularly important during alkaline stress. At elevated pH, Mn(II) is oxidized to Mn(IV)^45–47^ and Mn^2+^ hydroxide can promote RNA and DNA cleavage. Alkaline pH also exasperates oxidative stress by increasing free radical oxygen species (ROS)^48^. Conversely, Mn^2+^ can mitigate oxidative stress by serving as a cofactor for enzymes such as superoxide dismutase that break down ROS^49,50^. In fact, intracellular Mn^2+^ increases ∼1.5-fold in *E. coli* at alkaline pH^17^. Additionally, during oxidative stress, Mn^2+^ can substitute for iron in certain enzymes to partially restore enzymatic activity while limiting damage to the enzymatic active site from oxidized iron^51,52^. However, excess free Mn^2+^ is toxic^53^, necessitating tight homeostatic control. A dual Mn^2+^/pH sensing riboswitch that tunes Mn^2+^ exporter expression may therefore provide a robust mechanism to balance protection and toxicity in ecological niches that impose alkaline stress.

The *yybP-ykoY* aptamer core harbors binding sites for both Mg^2+^ and Mn^2+^, underscoring a likely role for Mg^2+^ in pre-organizing docking. Indeed, prior smFRET studies of *yybP-ykoY* aptamers observed no docked traces in monovalent cation-only buffers (K^+18,22^ or Na^+21^): without divalents, these aptamers were completely undocked (SU), whereas near-physiological Mg^2+^ promoted both DU and DD conformations in which the two helical legs transiently dock^18,21,22^. Consistent with these observations, Mg^2+^ alone is sufficient for *alx* to sample an SD conformation at pH 7.2, which may reduce the dynamic range for a Mn^2+^-dependent response at neutral compared to alkaline pH. Similar Mg^2+^-induced “folded-like” intermediates have been observed for other riboswitches, including preQ_1_ (where ligand binding subsequently locks the fully folded state)^54,55^, a nickel/cobalt (NiCo) riboswitch (where Mg^2+^ and Na^+^ promote folding and slow unfolding, even without ligand)^56^, and a fluoride riboswitch (where Mg^2+^ pre-folds the aptamer and coordinates with the ligand to lock the folded state)^57^. Together with our findings, these studies support a general model in which Mg^2+^ facilitates aptamer pre-organization, while ligand binding provides the final stabilization of the regulatory fold.

A central mechanistic theme is that the protonation state of RNA nucleotides can affect structure and catalytic activity^27,58^. Although the solution pK_a_ values for guanosine and uridine are near 9.2, and those for adenosine and cytidine are ∼3.5 and ∼4.2, respectively^27^, local RNA structural context can shift pK_a_ values toward physiological pH^59^. Our MD simulations suggest that protonation of an adenosine within the strained L2 capping loop can enable alternative base pairing of C19·A114^+^ (Fig. 4d) accompanied by formation of U18·A115, reorganizing the top helix of the aptamer’s “right” docking leg. We propose that this alternative pairing stabilizes a 4-nt L2 loop (Fig. 4b), thereby altering leg docking and Mn^2+^ binding in a pH-dependent manner. Because the docking timescales observed by smFRET are not readily accessible with direct simulation, it will be informative in future work to apply enhanced-sampling approaches for testing how adenosine protonation reshapes the docking landscape. More broadly, protonation-dependent non-canonical base pairing and conformational switching are established mechanisms in RNA structure and function^60,61^, including pH-tuned wobble formation that modulates microRNA processing through a G·A^+^ pair^62^, VS ribozyme activity via a C·A^+^ pair^28^, bacterial group II intron splicing based on a G·A^+^ pair^63^, eukaryotic spliceosomal catalysis enabled by a C·A^+^ pair^29,64^, as well as protonation-linked conformational switches that reprogram translation readthrough^65^ and frameshifting^65,66^. Our study expands this repertoire with a regulatory example in which pH reshapes RNA dynamics to allosterically modulate ligand binding and, consequently, gene expression.

Functionally, our *in vivo* data support a key role for L2 in pH-dependent translation activation in response to Mn^2+^ binding. Mutations in L2 perturb alkaline activation, with a strongly disrupting transversion (A114C) abolishing Mn^2+^ responsiveness and markedly attenuating pH- dependent activation (Fig. 4c,d). More modest L2 mutations reduce pH-dependent activation more prominently at slightly less alkaline pH (Extended Data Fig. 4e), indicating that additional RNA features contribute to the overall pH dependence. Consistent with this notion, prior co-transcriptional DMS probing data for *alx* showed that the absolutely conserved L3 adenosine that directly coordinates Mn^2+^ (A37) becomes substantially less reactive at alkaline pH in *alx*, whereas the corresponding position in the non–pH-responsive *mntP* does not^24^. In the current work, we swapped the L3 sequences and found that the native *alx* L3 is required for alkaline activation in cells: replacing it with the *mntP* L3 preserves Mn^2+^ responsiveness but eliminates pH-dependent activation (Fig. 5d), whereas inserting the *alx* L3 into *mntP* disrupts Mn^2+^ responsiveness. Finally, our phylogenetic analysis showed that both the *alx* L3 sequence and *alx*-like 3WJ architecture are conserved in *Bacillales* from alkaline-associated environments (Extended Data Fig. 10b), suggesting that allosteric coupling of binding-site sequence and global fold may be conserved to support growth under alkaline stress.

Beyond its physiological role, the *alx* aptamer also offers a design principle for synthetic RNA sensors. RNA aptamers inspired by natural riboswitches have long been pursued as ligand biosensors and fluorogenic imaging tools^67–69^, and recent efforts have sought single-molecule RNA sensors that integrate multiple inputs^70^. Such compact RNA sensors could complement established protein circuit strategies that require multiple interacting components^71,72^. Natural multi-input control is often achieved by tandem riboswitches, in which multiple aptamers act as sequential checkpoints—either amplifying output by sensing the same ligand or implementing combinatorial control with distinct ligands^73–75^. While effective *in vivo*, tandem riboswitch architectures require separate, compatible ligand-binding domains. In contrast, we show that *alx* represents a minimal, two-input allosteric sensor that readily integrates two distinct signals—pH and Mn^2+^—using only a single aptamer, providing a potential blueprint for engineering simpler multi-input RNA regulators.

## METHODS

### Protein

Q5 High-Fidelity DNA Polymerase, T4 DNA Ligase, PstI-HF, and BamHI-HF were purchased from New England Biolabs. Glucose oxidase and catalase were purchased from Sigma Aldrich and resuspended according to manufacturer instructions. Details and catalog numbers for all commercially available proteins are available in Supplementary Table 1.

### Oligonucleotide design

Custom oligos for smFRET were ordered from Revvity Inc (formerly Horizon Discovery). Custom oligo modifications of 5ʹ-Cy3, 5ʹ-Cy5, and 3ʹ-biotin were added where indicated (Supplementary Table 2). All custom oligos were desalted, deprotected, and purified via HPLC. Other oligos used in this study were ordered from Eton Bioscience Inc and purified via desalting. All oligos used in this study are tabulated in Supplementary Table 2.

### Bacterial strains

The strain used in this study is a derivative of *E. coli* K-12 (MC4100). The Δ*alx*::Kan mutation was sourced from the Keio collection^76^ and introduced into the *E. coli* MC4100 strain via P1 transduction. All strains used in this study are tabulated in Supplementary Table 3.

### Plasmids

The plasmid carrying the *alx* translation *lacZ* reporter gene fusion under the control of a native *E. coli* promoter is pRA54 and the empty vector plasmid is pMU2386. These were constructed in prior work and used as controls in this work (Supplementary Table 4)^17^. Translation *lacZ* reporter gene fusions of *alx* mutants were constructed via sequential PCR amplification from pRA54. The first amplification of pRA54 used forward primer 0504, which contains a PstI restriction site, and a reverse primer appropriate for introducing the desired L2 or L3 mutation(s) (Supplementary Table 2). The second amplification of pRA54 used a corresponding forward primer to introduce the desired L2 or L3 mutation(s) and reverse primer 0505, which contains a BamHI restriction site (Supplementary Table 2). The final PCR used primers 0504 and 0505 to combine these two fragments into the full mutant *alx* translational fusion insert. Translation *lacZ* reporter gene fusions of *mntP* mutants were constructed via sequential PCR from pRA57. The first amplification of pRA57 used forward primer 0506, which contains a PstI restriction site, and reverse primer 0511 appropriate for introducing the desired L3 mutations (Supplementary Table 2). The second amplification of pRA57 used corresponding forward primer 0510 to introduce the desired L3 mutations and reverse primer 0507, which contains a SalI restriction site (Supplementary Table 2). The final PCR used primers 0506 and 0507 to combine these two fragments into the full mutant *mntP* translation fusion insert. Then, the *alx* translational fusions were cloned into the PstI and BamHI sites of the single copy plasmid pMU2386. The *mntP* translation fusions were cloned into the PstI and SalI sites of the single copy plasmid pMU2386. The plasmids were assembled via restriction-ligation using T4 DNA Ligase (New England Biolabs) and sequence-verified by Sanger sequencing. All plasmids used in this study are tabulated in Supplementary Table 4.

### smFRET data acquisition

smFRET experiments were performed using the Oxford Nanoimager (ONI) microscope in total internal reflection fluorescence (TIRF) mode. Movies were collected at 22 °C for 1,200 frames with 100 ms exposure time. PEG-passivated quartz slides were prepared as described in previous work^77^. The surface of the slide was incubated with 0.2 mg/mL streptavidin for 10 minutes, then washed three times with the appropriate 1X smFRET buffer (50 mM HEPES pH 7.2 or pH 8.5, 100 mM KCl). RNA oligos were hybridized in a 1:20 ratio (10 nM alx_apt_oligo_2_Cy5_Bio:200 nM alx_apt_oligo_1_Cy3) (Supplementary Table 2), heat denatured at 90 °C for 2 min, snap cooled on ice for 3 min, and refolded in 1X folding buffer (50 mM HEPES pH 7.5, 100 mM KCl, 1 mM MgCl_2_) at room temperature for 15 min. Diluted hybridized RNA samples (10-30 pM relative to alx_apt_oligo_2_Cy5_Bio) were immediately applied to the slide, incubated for 5 min at room temperature, then washed three times with appropriate 1X smFRET buffer (50 mM HEPES pH 7.2 or pH 8.5, 100 mM KCl). A freshly prepared enzymatic oxygen scavenging system (1X smFRET buffer, 5 mM MgCl_2_, 44 mM glucose, 165 U/mL glucose oxidase from *Aspergillus niger*, 5 mM Trolox, 2,170 U/mL catalase from *Corynebacterium glutamicum*) was added along with MnCl_2_, where indicated. Raw movies were collected for 120 sec with direct green laser (532 nm, 18% power) excitation for 100 s to detect smFRET, followed by addition of direct red laser (640 nm, 8% power) excitation for the final 20 s to identify the presence of Cy5.

### smFRET data analysis

Location of molecules and their corresponding fluorophore intensity traces were extracted from the raw movie files (see above) using custom MATLAB codes^22^. Traces were visualized and manually selected using MATLAB based on the following criteria: minimum intensity for Cy3 (200 A.U.), single step photobleaching of the dyes, and either longer than 20 s or having ≥1 clear transition between low and high FRET states. Selected traces were then background-subtracted to correct and minimal bleed through. The FRET ratio was calculated as I_A_/I_A_+I_D_ where I_A_ and I_D_ are the background-corrected intensities of Cy5 (acceptor) and Cy3 (donor), respectively. FRET histograms were made using the first 10 frames of all traces in each condition and were fit with a sum of Gaussians using OriginPro 10.2. For kinetic analysis, traces were idealized using a two-state model corresponding to low FRET (undocked) and high FRET (docked) states using the segmental k-means algorithm in QuB software as previously described^25,78^. Cumulative dwell time histograms were plotted from all extracted dwell times and fit with double exponential functions using OriginPro 10.2 to obtain lifetimes in undocked (t_undock_) and docked (t_dock_) states. The rate constants for docking and undocking were calculated as follows: k_dock_ = 1/t_undock_ and k_undock_ = 1/t_dock_. Kinetics were calculated using both the short and long dwell times to calculate the fast and slow rate constants, respectively. Idealized smFRET traces were used for creating transition occupancy density plots (TODPs), which show the fraction of traces molecules undergo ≥1 transition between FRET states. In TODPs, dynamic traces showing a FRET transition (regardless of the number of transitions in the trace) and static traces (with no transitions in the trace) are weighted equally to avoid overrepresenting traces with many fast transitions. Graphs were visualized in GraphPad Prism.

### β-galactosidase assays

Overnight culture of each strain was diluted 1:100 into LBK (LB media where KCl replaces NaCl) with appropriate antibiotic at either pH 6.8 or pH 8.5, with or without 1 mM supplemented MnCl_2_, where indicated. The cultures were grown to mid-log phase and β-galactosidase assays were performed using the Miller method^79^. Briefly, bacterial growth was measured using OD_600_ before cultures were placed on ice. 100 µL cells were lysed via vortexing in 900 µL sodium phosphate buffer (100 mM sodium phosphate pH 7.0, 10 mM KCl, 2 mM MgSO_4_, 38 mM 2-mercaptoethanol), 20 µL 0.1% SDS, and 20 µL chloroform. Substrate *O*-nitrophenyl- b -D-galactopyranoside (ONPG) was added and the reaction was stopped with 1 M Na_2_CO_3_ when a yellow color appeared, or after 35 min. After centrifuging 5 min at 16,000*g*, absorbance was measured at 420 nm (A_420_) to calculate activity using Miller units. Each culture was tested in duplicate and A_420_ values were averaged. Each final reported value is an average of three replicates with standard deviation. Statistical analysis was done in GraphPad Prism using a two-way ANOVA with Tukey post-hoc test for multiple comparisons.

### Sequence alignment and phylogenetic analysis

Sequences for *yybP-ykoY* riboswitches were downloaded from Rfam (Rfam number: RF00080)^34^ and aligned using Kalign multisequence alignment (https://www.ebi.ac.uk/jdispatcher/msa/kalign). The sequence alignment was visualized using Jalview^80^. The L3 consensus sequence was visualized and exported from Jalview. The sequences were filtered by the *alx* L3 sequence and *alx* L3-containing sequences were exported (Supplementary Table 7). The phylogenetic tree was generated using phyloT (https://phylot.biobyte.de/) and the original Rfam organism names were updated according to NCBI taxonomy^81^, where indicated in bold (Supplementary Table 7). The phylogenetic tree was visualized using Interactive Tree Of Life (iTOL, https://itol.embl.de/).

### Structure prediction and modeling

For *yybP-ykoY* riboswitch sequences containing *alx* L3, structure predictions were generated using RNAfold with default settings^35^ and were manually noted as having *alx* or *mntP*-like architecture (Supplementary Table 7). Two-dimensional RNA structure models were visualized in VARNA^82^. Crystal structures of the Mn-bound *alx* aptamer (PDB: 9JGM)^19^ and the Mn-bound *X. oryzae* aptamer (PDB: 6N2V)^18^ were visualized and colored in ChimeraX^83^. Crystal structure alignments were also done in ChimeraX using the default matchmaker settings.

### Co-transcriptional SHAPE-MaP single-nucleotide trajectories

Previous published raw sequencing data from co-transcriptional DMS-MaP of the *alx* riboswitch is available in the SRA under project number PRJNA1197522 (https://www.ncbi.nlm.nih.gov/sra)^24^. This co-transcriptional structure probing was performed *in vitro* using the Variable Length Transcription Elongation Complex RNA Structure Probing (TECprobe-VL) method^84^. This data from prior work contained the reactivities with DMS for nucleotides of interest at pH 7.2, pH 8.5, and ±Mn at every RNA transcript length. In the current work, we plotted the single nucleotide trajectories for nucleotides of interest in GraphPad Prism.

### Molecular dynamics simulations

#### System setup

Molecular dynamics (MD) simulations were performed to characterize the conformational dynamics of the *alx* riboswitch aptamer under different structural and protonation conditions. Four simulation setups were prepared for standard MD conditions, combining two starting conformations (native and C19·A114 wobble base-pair states, see below for the procedure used to model the wobble pair) with two protonation schemes for residue A114: fixed protonated and fixed deprotonated.

Stochastic titration constant-pH MD (CpHMD) simulations were performed using a discrete protonation framework as described in other works^33,85,86^. CpHMD simulations were prepared starting from either the native and wobble states, with a single titrable residue (A114). In all simulations, all other nucleotides were kept in their standard neutral protonation states.

All systems were derived from each chain (A-D) of the Mn-bound *alx* aptamer crystal structure (PDB: 9JGM). Structural divalent ions present in the binding pocket were retained, though the Mn²⁺ ions were replaced by Mg²⁺ due to the lack of reliable force-field parameters for Mn²⁺. No additional divalent ions were included.

Each system was solvated in a dodecahedral box to physiological conditions using 38,642 water molecules, 130 K⁺ ions, and 31 Cl⁻ ions, to match a salt concentration of 100 mM and ensuring charge neutrality. System topologies were generated using a modified XOL3 force field for RNA^33,87^, Joung–Cheatham parameters for monovalent ions^88^, and the microMg model for Mg²⁺ ions^89^. Water molecules were described using the OPC model^90^.

All simulations were performed using GROMACS 2022.3^91^ coupled to PLUMED 2.8.2^92,93^ at constant temperature (310 K) using a v-rescale thermostat^94^, while the pressure (1 bar) was maintained using a c-rescale barostat^95^. Covalent bonds were constrained using the LINCS algorithm^96^, and long-range electrostatic interactions (at distance larger than 1.0 nm) were treated using the particle-mesh Ewald (PME) method^97^. Each system was minimized before equilibration and other protocols.

#### Generation of wobble base-pair conformations

Wobble base-pair starting structures were generated from the native conformation for each crystallographic chain (A–D). The native U18·A114 base pair was converted to an C19·A114⁺ wobble base pair using eRMSD-based steered MD simulations^32,33^. Steering enforced the formation of C19·A114 and U18·A115 base pairs. The latter was found pivotal to the wobble base pair stability, as found in hairpin structures studied by Wilcox and Bevilacqua et al^30^. One of these sequences was used to generate a target geometry.

Steering protocols were performed using the eRMSD metric as implemented in PLUMED 2.8.2., which accounts for nucleobase distances and relative orientation to measure structural similarity. The eRMSD was computed using nucleobases 18, 19, 114, and 115. A harmonic moving restraint with force constant κ = 10 000 kJ mol⁻¹ was applied to drive the eRMSD from its native value toward 0, corresponding to the target wobble base pair geometry of the reference structure. Each steered MD simulation was run for 200 ns.

For each chain (A-D), two replicates were performed corresponding to fixed protonated and fixed deprotonated states of A114 (see Supplemental Information). Following the steering phase, a second equilibration run of 200 ns was performed while preserving the same eRMSD restraint parameters to allow relaxation of the remaining RNA structure. An additional unrestrained equilibration run of 1 µs was then performed prior to production simulations, which are described in the following section. Wobble base-pair formation was verified through eRMSD convergence and analysis of hydrogen-bond distances over simulation time.

#### Production simulations

Production simulations were carried out at 310 K and 0.1 M KCl using either standard MD or CpHMD, depending on the protonation scheme. For systems with fixed protonation states, two independent 1 µs MD replicates were performed for each crystallographic chain (A-D), starting from either the native or wobble base-pair conformations, except for the native protonated and wobble deprotonated schemes that were structurally unstable (Extended Data Fig. 5a & 6a).

For CpHMD simulations, eRMSD restraints were applied to preserve the structural stability of native and wobble states throughout the simulations. Two independent replicates were performed for each conformational state and each chain (A-D) across distinct pH ranges: pH 7.0–12.0 for the wobble state and pH 3.0–7.0 for the native state. The pH titration range for the wobble state was selected based on changes in activity observed at pH 8.5 in the experimental assays. The pH titration range for the native state was chosen considering that the pKa of a Watson-Crick base pair is lower than the aqueous pKa of a free adenine (3.5)^30,31^.

#### Trajectory analysis

Trajectory analyses were performed to characterize the conformational stability, protonation behaviour of A114, L2 loop length, and key hydrogen-bond interactions of the *alx* riboswitch across all production simulations. Structural convergence and stability over time of native and wobble conformational states was quantified using the eRMSD metric. For CpHMD simulations, residue-specific protonation time-series were verified, while averages were computed by averaging the protonation state of A114 of each simulation at each pH values. Titration curves were generated from these averages and fit to the Henderson-Hasselbalch equation to estimate the microscopic pKa values. Hydrogen-bond interactions were identified using geometric criteria on donor-acceptor distances (<0.35 nm) and angles (>150°) analyzed with MDanalysis. L2 loop length was determined indirectly by the presence of the C19·G23 base pair, or its absence coupled with the C19·A114⁺ wobble base pair. Statistical uncertainties for all reported observables were estimated using a block bootstrap approach applied to the full production trajectories, where time series were partitioned into 10 blocks each trajectory and resampled to obtain mean values and standard errors.

## DATA AVAILABILITY

The modified nucleic acid parameters used in the st-CpHMD are available at https://github.com/Tomfersil/CpH-MetaD. All the input scripts are available at https://github.com/Tomfersil/Alx-Riboswitch-MD/. PLUMED input files are also available on PLUMED-NEST^92^ with plumID:26.001. Processed trajectories are available on Zenodo at https://doi.org/10.5281/zenodo.18462847. Source data for single molecule FRET assays are available through the University of Michigan DeepBlue deposit (https.doi.XXXXX, TBD) and will be accessible upon manuscript acceptance.

## ACKNOWLEDGEMENTS

We thank C. Stephen for helpful initial conceptualization and thank all Mishanina lab members for their valuable feedback. This work was supported by the National Institutes of Health (NIGMS ESI grant R35GM142785 to T.V.M. and MIRA grant R35 GM131922 to N.G.W.), the Italian National Centre for HPC, Big Data, and Quantum Computing (CN00000013 to G.B.), Next Generation EU initiative through PRIN 2022 (2022Z4FZE9 to G.B. and T.F.D.S.), European Molecular Biology Organization, EMBO (ALTF-399/2022 to T.F.D.S.), the National Science Foundation (MCB 2516225 to T.V.M. and MCB 2140320 to N.G.W.), and the National Science Foundation Graduate Research Fellowship Program (DGE-2038238 to D.P.). Any opinions, findings, and conclusions or recommendations expressed in this material are those of the author(s) and do not necessarily reflect the views of the National Science Foundation.

## AUTHORS AND AFFILIATIONS

**Department of Chemistry and Biochemistry, University of California, San Diego, 9500 Gilman Dr, La Jolla, CA, USA**

Danea Palmer, Avery Ontiveros, & Tatiana V. Mishanina

**Single Molecule Analysis Group and Center for RNA Biomedicine, Department of Chemistry, University of Michigan, Ann Arbor, MI, USA**

Adrien Chauvier & Nils G. Walter

**Scuola Internazionale Superiore di Studi Avanzati, via Bonomea 265, Trieste, 34136, Italy**

Tomás F. D. Silva & Giovanni Bussi

## Contributions

Conceptualization: D.P., T.V.M., N.G.W, and G.B. Experimental design: D.P., N.G.W., A.C., T.F.D.S., and G.B. Investigation: A.C. and D.P. (smFRET experiments and data analysis); D.P. and A.O. (*in vivo* experiments and data analysis); T.F.D.S. (molecular dynamics simulations and data analysis). Writing (original draft): D.P. and T.V.M. Writing (review and editing): N.G.W., A.C., T.F.D.S., G.B., D.P., and T.V.M. Supervision: T.V.M., N.G.W., and G.B. Funding acquisition: T.V.M., N.G.W., and G.B.

## Corresponding Author

Correspondence to Nils G. Walter and Tatiana V. Mishanina

