## Extended Data for "pH-dependent allosteric remodeling of a bacterial riboswitch couples alkaline activation to metal sensing"

**a**

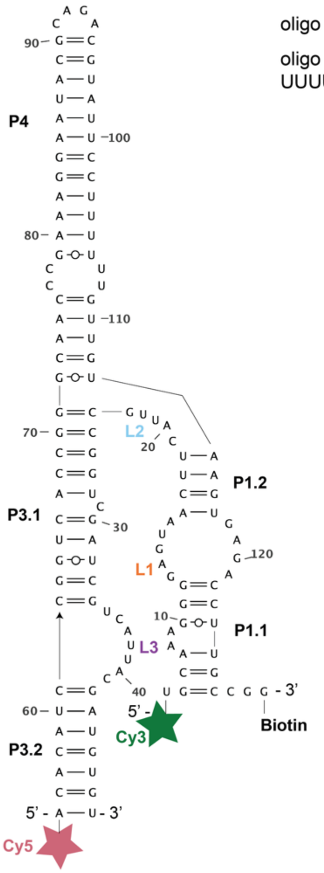


**b**

oligo 1: 5'-**Cy3**-UGCAAAGGGGAGUAACUUCAUUGCCGGUCGAUCGUCAUUACGAUGUGU-3'

oligo 2: 5'-**Cy5**-ACACAUCCGGUCACCGGGCAACCCGAAAGGAAUACGCAGACGUAUUCCU UUUUUGUUGUAAGUGAGACCUUGCCGG-**Biotin**-3'

**Extended Data Fig. 1: smFRET construct design.**

**a,** Sequence and secondary structure for the *alx* aptamer construct used for smFRET. Positions of Cy5, Cy3, and biotin labels are displayed on the secondary structure. **b,** The oligo sequence and modifications ordered for smFRET experiments.

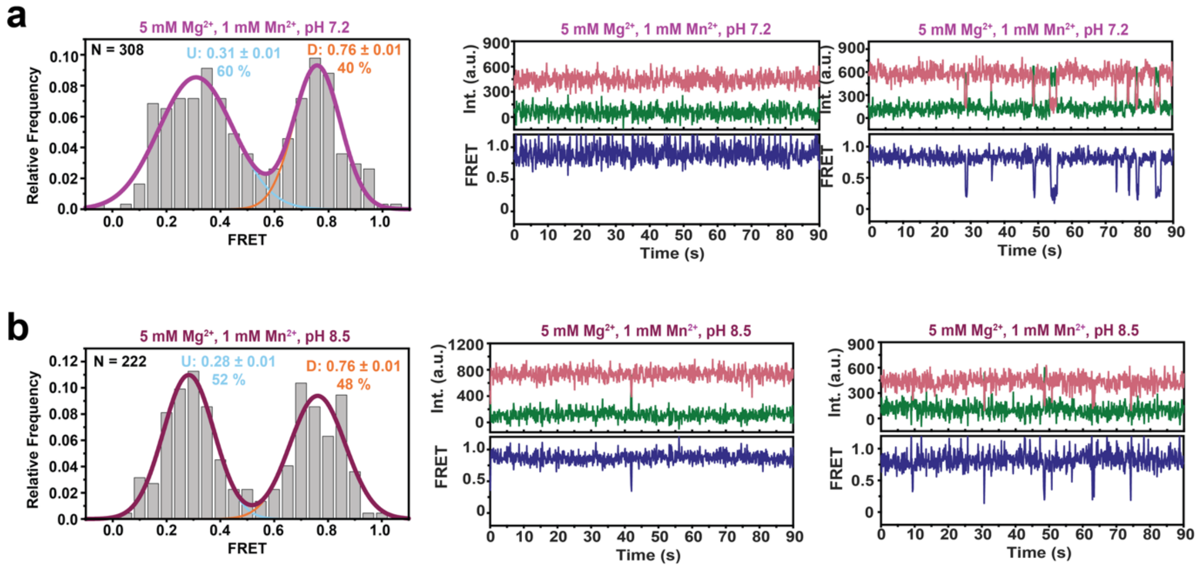


**Extended Data Fig. 2: *alx* aptamer docking at saturating [$Mn^{2+}$] and neutral vs alkaline pH.**

**a,b,** Histograms and representative traces showing the fraction of low and high FRET molecules with 1 mM $Mn^{2+}$ at **(a)** pH 7.2 and **(b)** pH 8.5. Histograms display the Gaussian peaks for low FRET, high FRET, and cumulative fit with the percentage of each FRET state and the number of molecules (N) analyzed.

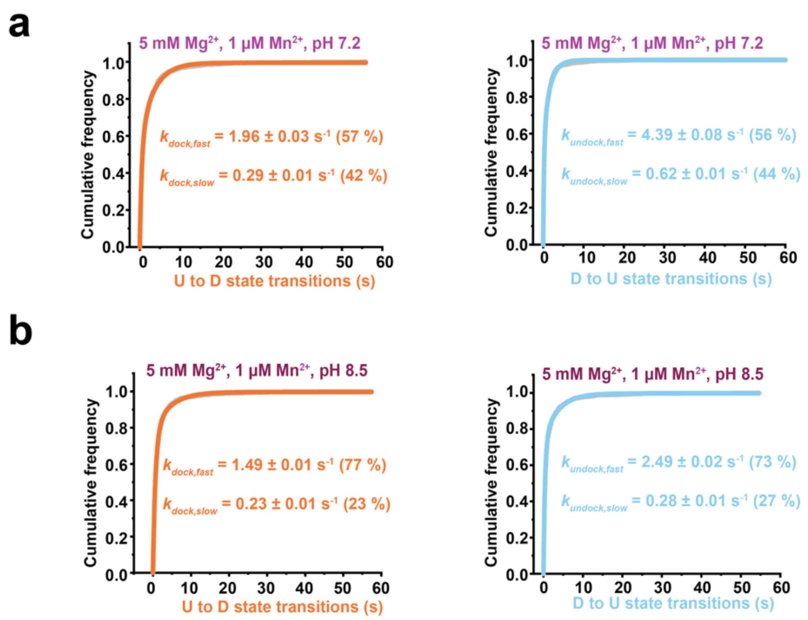


**Extended Data Fig. 3: *alx* aptamer docking kinetics at 1 µM $Mn^{2+}$ and neutral vs alkaline pH.**

**a,b,** Kinetics for docking (U to D state transitions) and undocking (D to U state transitions) based on the cumulative dwell times in each state fit with double-exponential functions for **(a)** pH 7.2 with 5 mM $Mg^{2+}$ and 1 µM $Mn^{2+}$ and **(b)** pH 8.5 with 5 mM $Mg^{2+}$ and 1 µM $Mn^{2+}$.

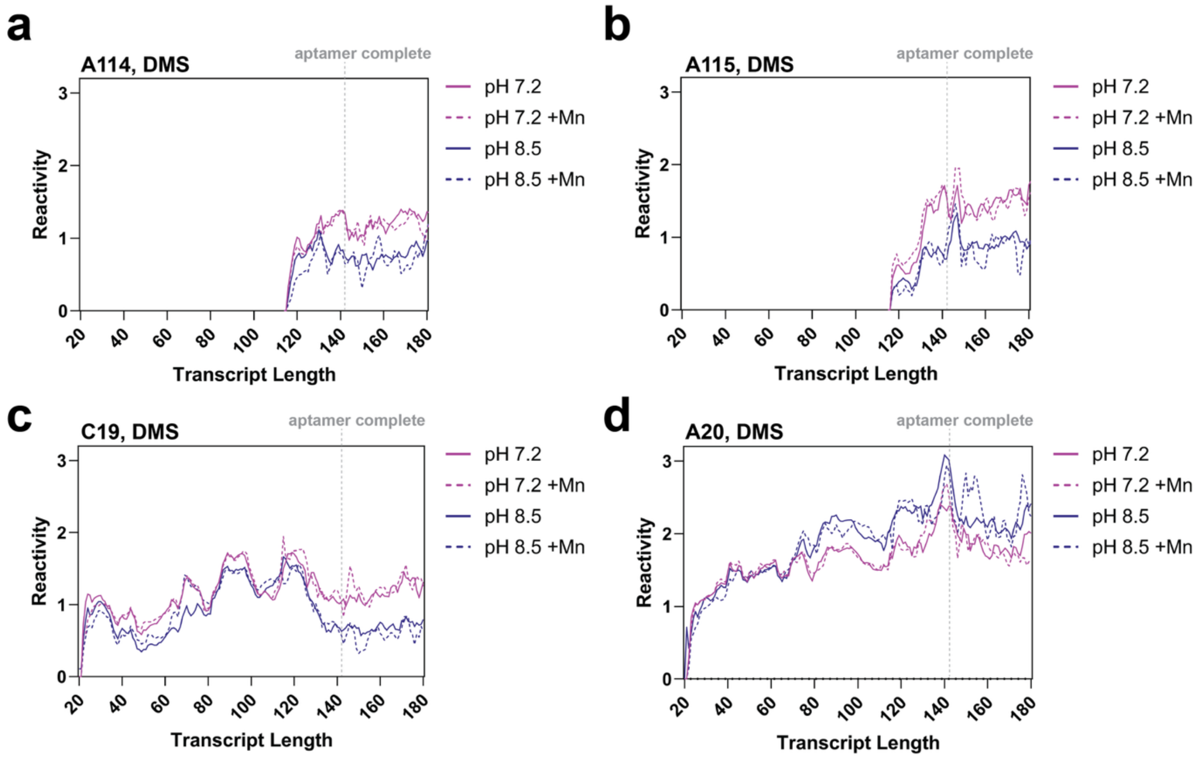


**Extended Data Fig. 4: Single-nucleotide DMS reactivity trajectories for L2 nucleotides.**

**a-d,** Single-nucleotide trajectories displaying transcript length-dependent reactivity changes for L2 nucleotides. DMS reactivity data from Stephen et al[24] plotted for pH 7.2 and pH 8.5 without $Mn^{2+}$ (solid lines) and with $Mn^{2+}$ (dashed lines) for the following nucleotides within the *alx* aptamer: **(a)** A114, **(b)** A115, **(c)** C19, and **(d)** A20.

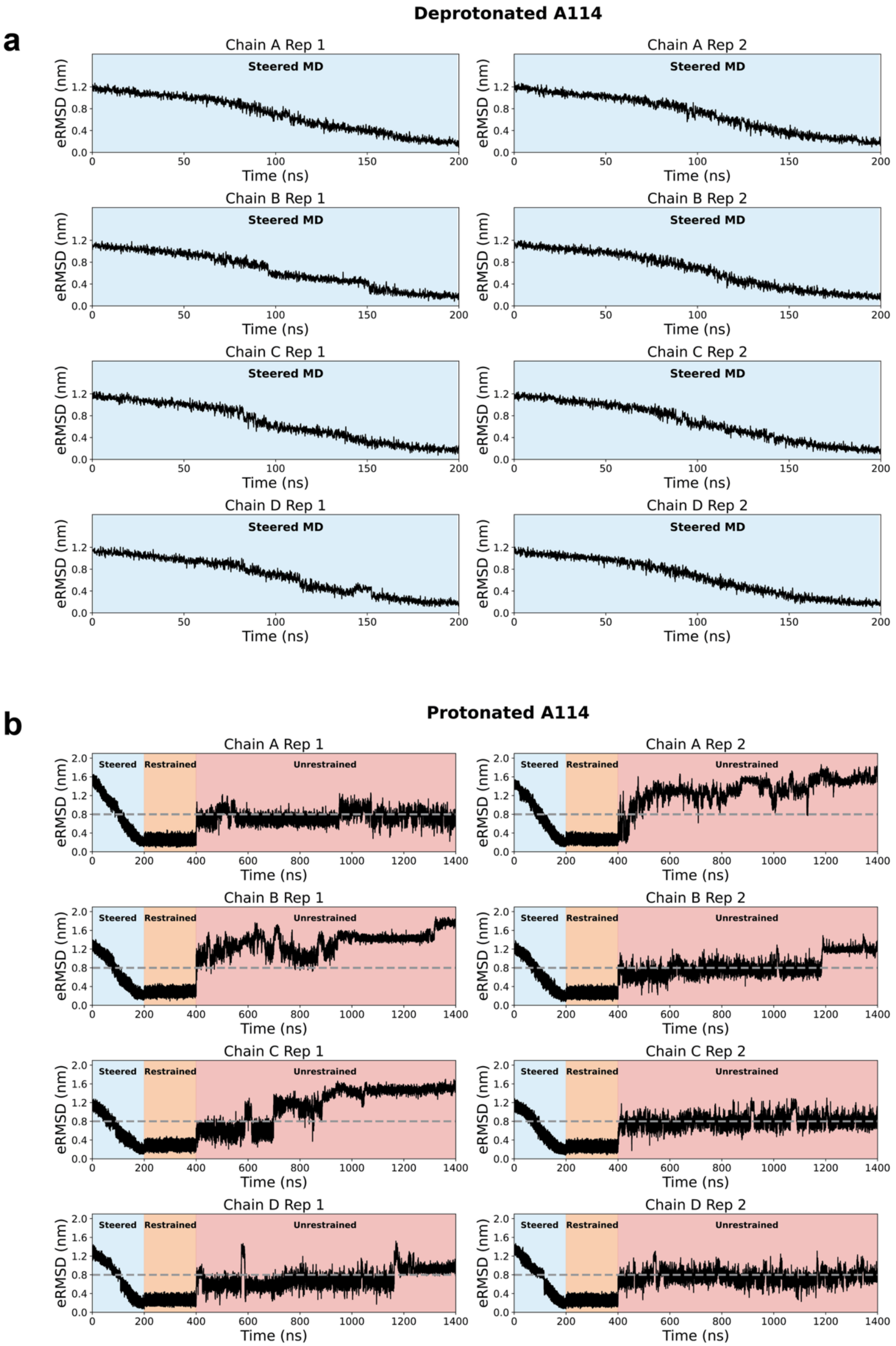


**Extended Data Fig. 5: Molecular dynamics protocol to generate wobble structure with a deprotonated and protonated A114.**

**a,b,** Time series of the eRMSD for each crystallographic chain (A–D) and replicate obtained from the steered, restrained, and unrestrained MD simulations of *alx* aptamer with a **(a)** deprotonated A114 or **(b)** protonated A114. During the steered step, the system is driven from the native crystal structure toward the target wobble reference geometry, forming the C19·A114 and U18·A115 base pairs. The restrained step allows relaxation of the remaining structure, while the unrestrained phase assesses the stability of the generated structure. Dashed lines indicate the eRMSD cutoff (0.8), below which conformations are considered similar to the reference structure. Restrained and unrestrained simulations for the deprotonated system are not performed since hydrogen bonds necessary to maintain the wobble state could not be formed.

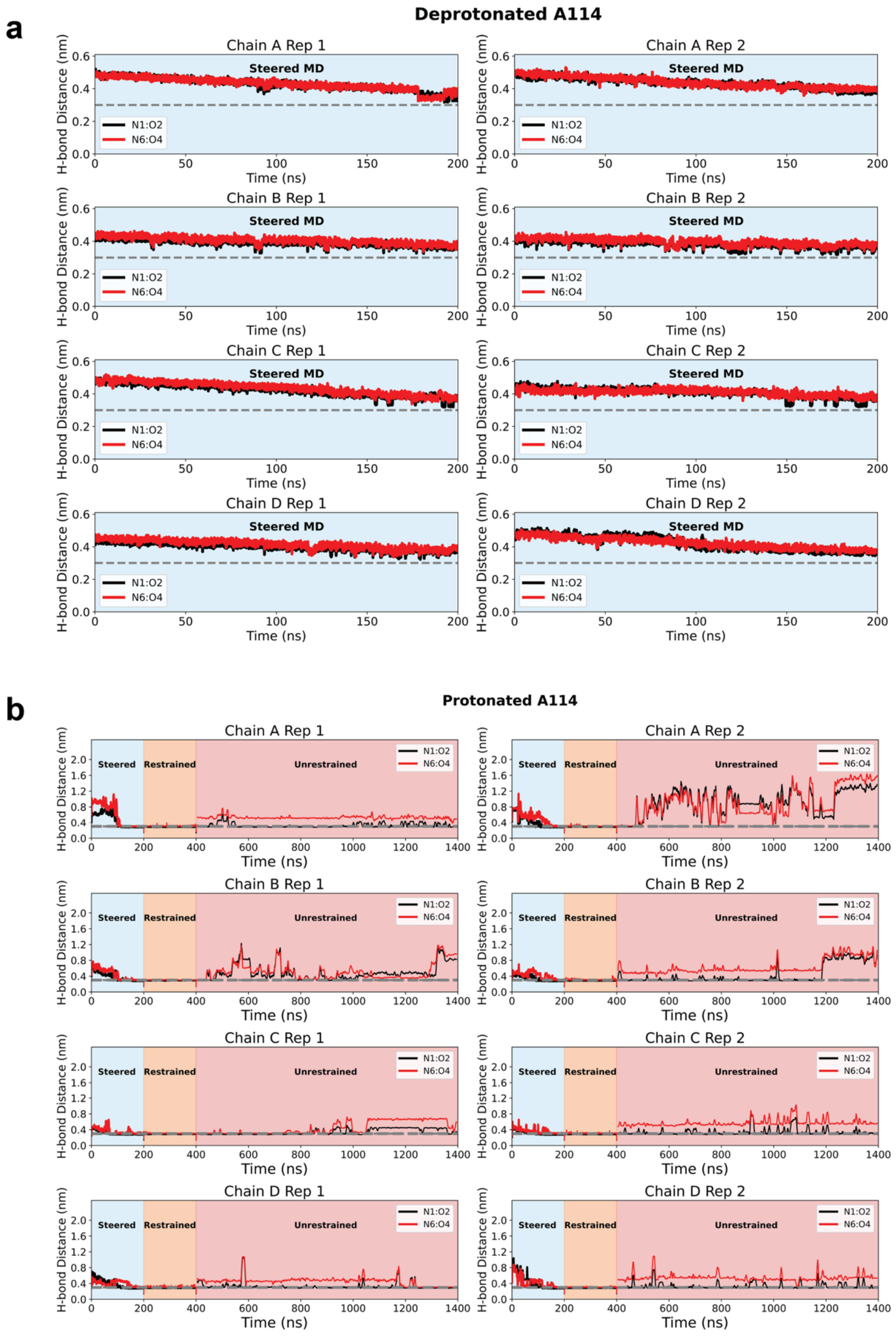


**Extended Data Fig. 6: Hydrogen-bond criterion for the formation of wobble C19·A114⁺.**

**a,b,** Time series of hydrogen-bond distances within the C19·A114 wobble base pair. Distances between the N1-O2 and N6-O4 atom pairs are shown for each crystallographic chain (A–D) and replicate during the steered MD simulations of the *alx* aptamer with **(a)** deprotonated A114 or **(b)** protonated A114. In (a), the measured hydrogen-bond distances fail to meet the distance criteria of a wobble base pair, supporting the data in Fig. 4 and 5. While in (b), conformational rearrangements, as shown by the large fluctuations, occur to facilitate convergence toward the hydrogen-bond distances found in the reference wobble structure. Dashed lines indicate the heavy-atom hydrogen-bond cutoff (0.3 nm), below which a stable hydrogen bond is considered formed.

**a**

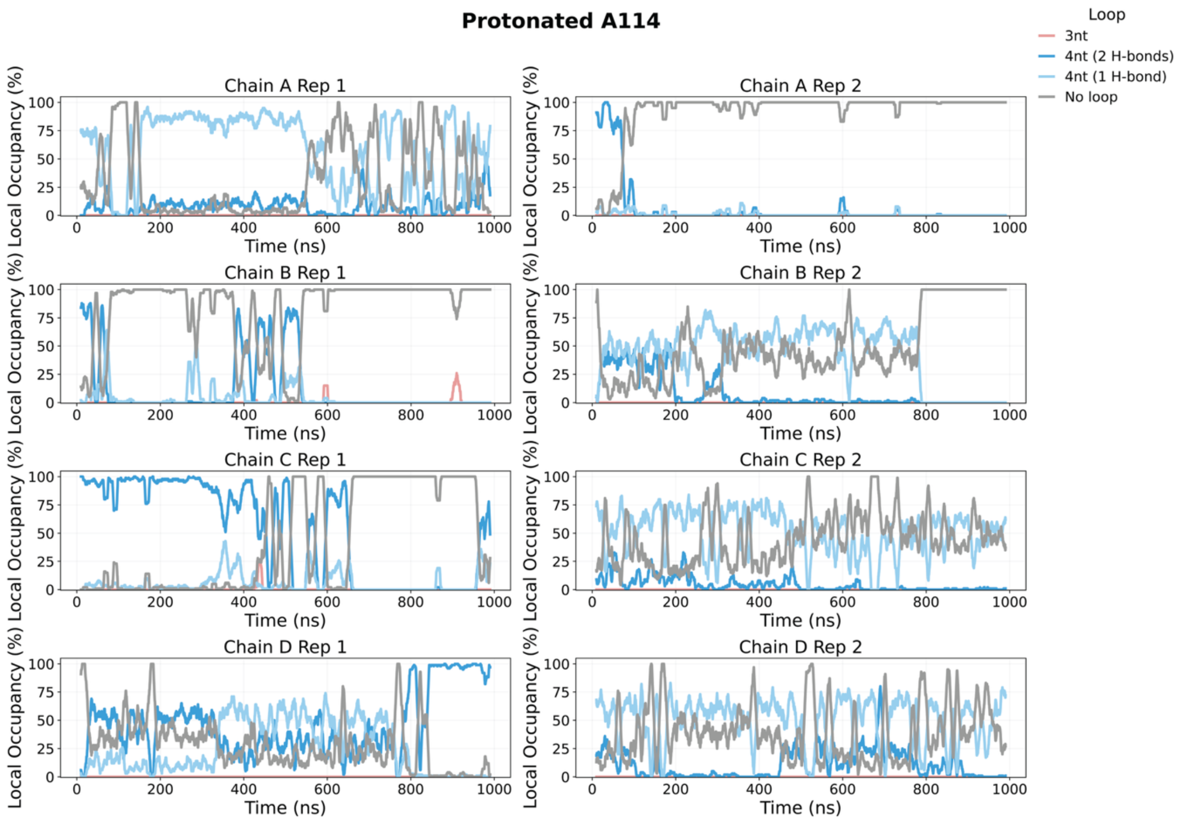
Protonated A114
Loop
3nt
4nt (2 H-bonds)
4nt (1 H-bond)
No loop
Chain A Rep 1
Chain A Rep 2
Chain B Rep 1
Chain B Rep 2
Chain C Rep 1
Chain C Rep 2
Chain D Rep 1
Chain D Rep 2
Local Occupancy (%)
Time (ns)

**b**

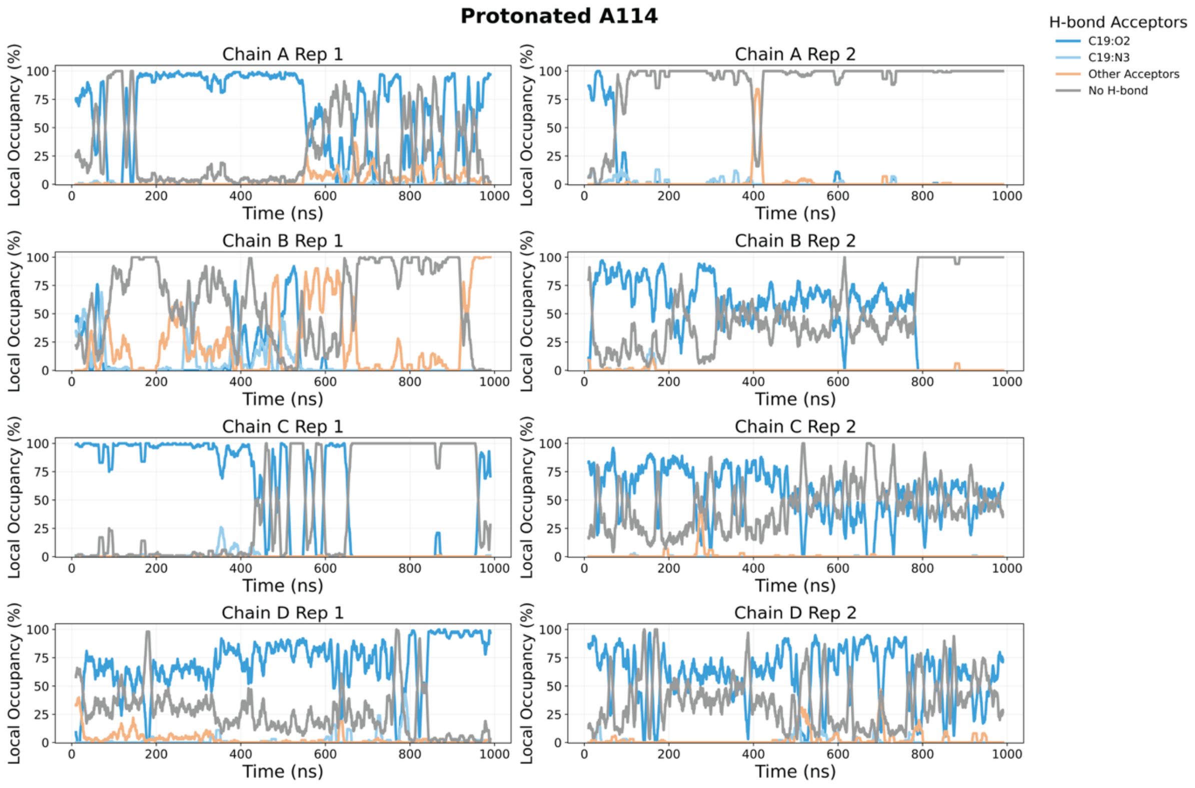
Protonated A114
H-bond Acceptors
C19:O2
C19:N3
Other Acceptors
No H-bond
Chain A Rep 1
Chain A Rep 2
Chain B Rep 1
Chain B Rep 2
Chain C Rep 1
Chain C Rep 2
Chain D Rep 1
Chain D Rep 2
Local Occupancy (%)
Time (ns)

**Extended Data Fig. 7: L2 loop length depends on the number of hydrogen bonds (H-bonds) and H-bond acceptor competition in the C19· A114$^+$ wobble structure.**

**a,** Relative occupancy of L2 loop-length states is defined by the number of H-bonds formed between C19·A114$^+$. Several transitions between multiple loop states are identified, whereas the 3-nt loop state is largely absent due to the lack of the stabilizing C19·G23 base pair found in the native structure. **b,** Time series of the relative occupancy of distinct H-bond acceptors interacting with the A114$^+$ N1H$^+$ donor. The A114$^+$ N1H$^+$ donor forms transient H-bonds with multiple acceptors, particularly the O2 and N3 of C19, thus preserving at least a partial wobble base pair. This partially stabilized state impacts the occupancy of L2 loop states and facilitates the observed transitions.  Time series plots were generated for each crystallographic chain (A–D) and replicate of the unrestrained MD simulations. Each time series was computed using a moving average window (n = 100 data points).

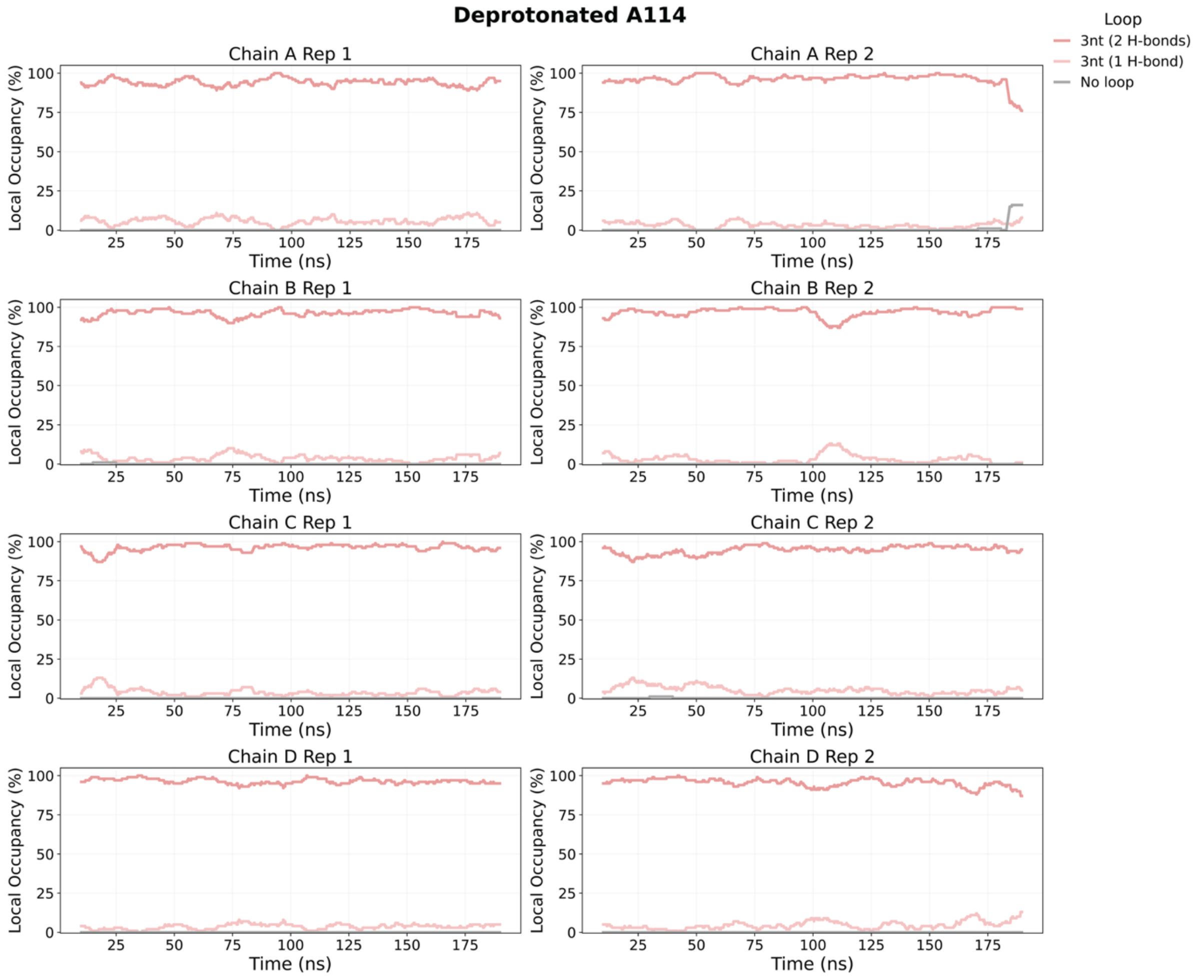


**Extended Data Fig. 8: L2 loop length is based on the number of hydrogen bonds (H-bonds) present in C19·A114 in the native structure.**

Time series of relative occupancy of L2 loop-length states defined by the number of H-bonds formed between C19·A114, for each crystallographic chain (A–D) and replicate of the unrestrained native MD simulations. Each time series was computed using a moving average window (n = 100 data points). No transitions are observed between loop states as the native structure is highly stable, particularly the C19·G23 and U18·A114 base pairs, in the absence of A114 protonation events to trigger conformational rearrangements.

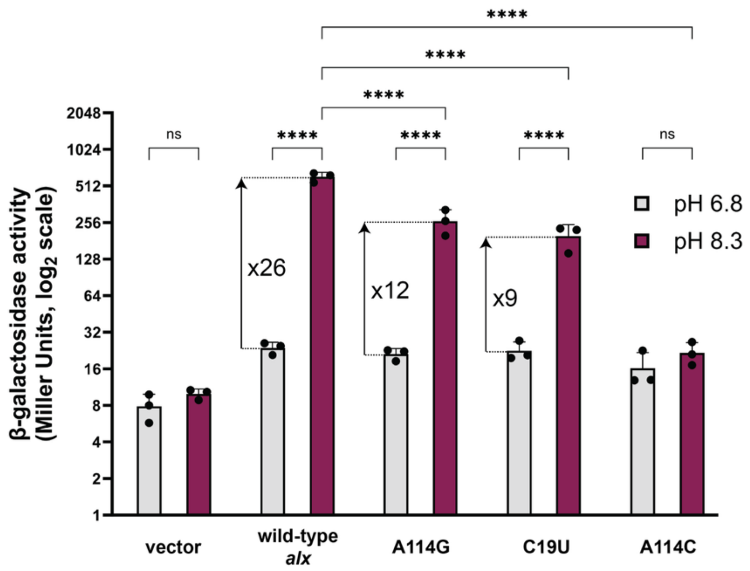


**Extended Data Fig. 9: L2 capping loop mutations reduce pH-dependent *alx* translation activation at pH 8.3**

Translational response of the L2 mutants compared to wild-type *alx* for growth media buffered at pH 8.3. Statistical analysis done using a two-way ANOVA with Tukey post-hoc test for multiple comparisons to assess significance ($^{ns}$ P>0.05, $^{*}$ P≤0.05, $^{**}$ P≤0.01, $^{***}$ P≤0.001, $^{****}$ P≤0.0001).

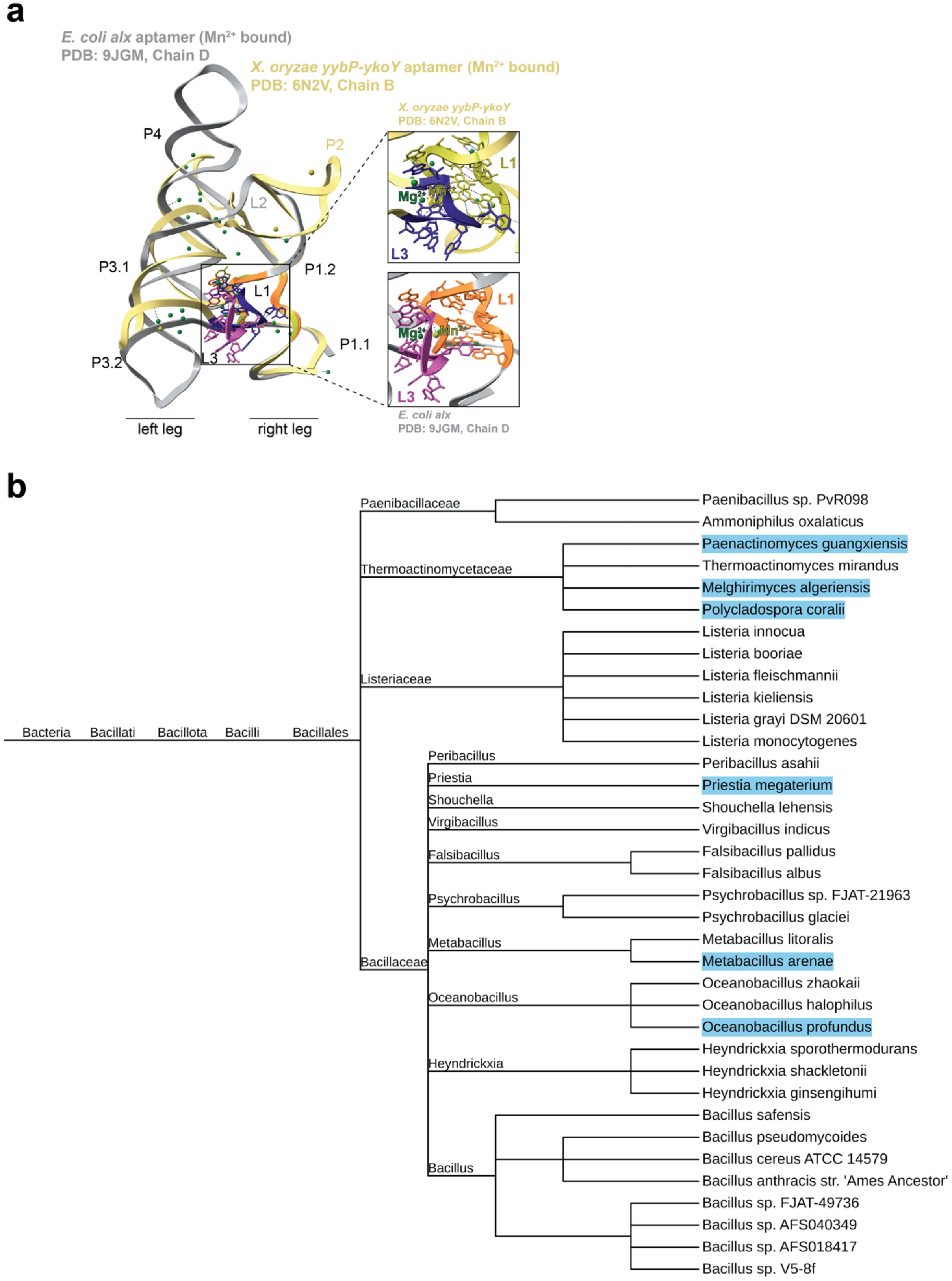


**Extended Data Fig. 10: L3 structure, sequence conservation, and phylogenetic analysis.**

**a,** Alignment of the *yybP-ykoY* family aptamer crystal structures for the *E. coli alx* (PDB: 9JGM, Chain D) and *X. oryzae* (PDB: 6N2V, Chain B). Individual aptamer $Mn^{2+}$ binding sites are displayed in the zoomed-in panels for both *alx* and *X. oryzae*. **b,** Phylogenetic tree of *yybP-ykoY* family aptamers (from Rfam) with the *alx* L3 sequence. The highlighted organisms have *alx*-like aptamer architecture.
